# TGF-β1 induced S100 family protein expression is associated with epithelial to mesenchymal transition states and poor survival in pancreatic cancer

**DOI:** 10.1101/2022.02.25.481888

**Authors:** Ronnie Ren Jie Low, Ka Yee Fung, Hugh Gao, Adele Preaudet, Laura F. Dagley, Jumana Yousef, Belinda Lee, Rune H. Larsen, Nadia J. Kershaw, Antony W. Burgess, Peter Gibbs, Frédéric Hollande, Michael D.W. Griffin, Sean M. Grimmond, Tracy L. Putoczki

## Abstract

Epithelial-mesenchymal transition (EMT) is a continuum that includes epithelial, partial EMT (P-EMT) and mesenchymal states, each of which are associated with cancer progression, invasive capabilities and ultimately metastasis. We have employed a lineage traced sporadic model of pancreatic cancer to generate a murine organoid biobank from primary and secondary tumors, including sublines that have undergone P-EMT and complete EMT (C-EMT). Using an unbiased proteomics approach, we found that the morphology of the organoids predicts the EMT state, with solid organoids associated with a P-EMT signature. We also observed that exogenous TGFβ1 induces a solid organoid morphology that is associated with changes in the S100 family, C-EMT and the formation of high-grade tumors. S100A4 may represent a useful biomarker to predict EMT state, disease progression and outcome for pancreatic cancer patients.

## INTRODUCTION

Pancreatic ductal adenocarcinoma (PDAC) is amongst the most lethal malignancies, with a 5-year survival of less than 12% (Siegel et al., 2020). Many patients succumb to the disease within the first 6 months, a reflection of late diagnosis, metastasis, and therapy resistance (Katz et al., 2009). A distinctive feature of PDAC is the progression of an early pancreatic intraepithelial neoplasia to advanced unresectable disease (Feldmann et al., 2007). During this process, the histopathological phenotype of a tumor changes from low grade, characterized by well-formed glandular epithelial structures, to high grade, whereby epithelial cells are diffuse throughout the desmoplastic stroma (Low et al., 2021; Schlitter et al., 2017). This process of dissemination has been linked to the activation of epithelial-mesenchymal transition (EMT), a cell-biological program that is essential to the early stages of embryogenesis, the formation of organs and exploited by cancer cells (Thiery, 2002; Yang et al., 2020). EMT supports pathological events associated with the loss of epithelial behaviour and the acquisition of mesenchymal features that enable invasion and metastasis (Thiery, 2002; Yang *et al*., 2020).

In PDAC, retrospective patient analysis, *in vitro* assays, xenograft models and genetically engineered mouse models (GEMMs) have suggested that a set of transcription factors (TF) enable EMT (Arumugam et al., 2009; Beerling et al., 2016; Chen et al., 2018; Krebs et al., 2017; Reichert et al., 2018; Rhim et al., 2012b; von Burstin et al., 2009; Wang et al., 2019; Zheng et al., 2015). However, we now know that the progression of PDAC is not solely driven by the classic EMT-TFs Snail or Twist (Zheng *et al*., 2015); although, the loss of the EMT-TF Zeb1 can result in the growth of a low grade tumor (Craene and Berx, 2013; Krebs *et al*., 2017; Lamouille et al., 2014). These EMT-TFs were originally considered part of a binary process, whereby cells transitioned into two distinct epithelial or mesenchymal states, the latter defined by the loss of expression an epithelial protein, E-Cadherin and the gain of expression of a mesenchymal protein, Vimentin (Yang *et al*., 2020). This concept has now shifted to reflect a continuum of partial EMT (P-EMT) states with overlapping transcriptional features that are found in multiple cancers, including PDAC (Aiello et al., 2018; da Silva-Diz et al., 2018; Hong et al., 2015; Huang et al., 2013; Jolly et al., 2018; Jolly et al., 2016; Jordan et al., 2011; Kalluri and Weinberg, 2009; Lovisa et al., 2015; Lovisa et al., 2016; Pastushenko et al., 2018; Williams et al., 2019; Yang *et al*., 2020). Cancer cells can also revert from P-EMT states to epithelial states through mesenchymal-epithelial transition (MET) (Beerling *et al*., 2016; Jolly et al., 2017; Reichert *et al*., 2018). This spectrum of heterogenous EMT states may be associated with differences in epithelial plasticity and cell migration, thereby influencing tumor progression, metastasis and response to treatment (Aiello *et al*., 2018; Lüönd et al., 2021). Targetable cell intrinsic mechanisms that drive either P-EMT or C-EMT remain elusive.

PDAC tumor cells reside within a dense desmoplastic stroma comprising cancer associated fibroblasts (CAFs) and associated extracellular matrix (ECM), which constitute more than 80% of the tumor mass (Dougan, 2017; Erkan et al., 2008). How the stroma modulates the behavior of tumor cells, and in particular the progression of EMT, is not clear. A primary focus has been placed on the cytokine TGFβ1, which is potently secreted by CAFs (Marzoq et al., 2019) resulting in autocrine signalling and differentiation into myofibroblastic CAFs that promote deposition of ECM proteins (Biffi et al., 2019; Choi et al., 2019; Shek et al., 2002). Paracrine TGFβ1 signalling can also act on benign neoplastic epithelial cells leading to cell cycle arrest or on neoplastic cells, to induce proliferation, motility and EMT (David et al., 2016; Gabitova-Cornell et al., 2020; Huang et al., 2020a; Huang et al., 2020b; Ligorio et al., 2019; Nicolás and Hill, 2003; Su et al., 2020; Zhang et al., 2014). How stromal derived and cell intrinsic mechanisms converge on EMT phentoypes in cancer cells is not clear, with the identification of transitions between the different stages of the EMT continuum technically challenging.

Here, we undertook an unbiased quantitative proteomics approach to explore the relationship between murine organoid morphology, the EMT continuum and tumor grade. We show that in a murine model of PDAC, organoids with a solid morphology have undergone P-EMT and form high grade tumors when transplanted into syngeneic hosts. We further explored the impact that contextual signals from cytokines derived from the stromal microenvironment had on the induction of EMT. We demonstrate that TGFβ1 can induce changes in organoid morphology and provide evidence that this is associated with the induction of C-EMT and alterations in S100 family expression. Collectively, our findings suggest that S100A4 may represent a useful biomarker to predict EMT state, disease progression and survival.

## RESULTS

### Murine organoids have two distinct morphological sub-types

The *Pdx*^Cre^; *Kras*^G12V^; *p53*^*R*172H^; *Rosa*^YFP^ (CKPY) GEMM of PDAC was utilised to generate a murine organoid (MO) biobank, as the YFP allele permitted identification (Figures S1A-B) and FACs isolation (Figure 1A) of metastatic lesions in the lung, liver and diaphragm (Table S1). Littermate *Pdx*^Cre^; *Rosa*^YFP^ (CY) mice were used to isolate epithelial cells from the ‘normal’ healthy pancreas (Figure 1A; Table S1). On average, ∼13% of the cells detected in CY pancreas were YFP+ epithelial cells, with no YFP+ cells detected in the liver, diaphragm or lung as expected (Figure S1C-D). For the CKPY mice, on average ∼17% of the cells detected in the pancreas were YFP+ epithelial cells, with YFP+ cells also detected in the liver, diaphragm, lung and blood, although the percentage of YFP+ cells detected in these organs varied between mice (Figure S1D). There was a 100% success rate for the generation of normal pancreatic MOs and pancreatic tumour MOs, while the success rates for generation of MOs from the metastatic lesions varied (Figures S2A-B).

**Figure 1.**
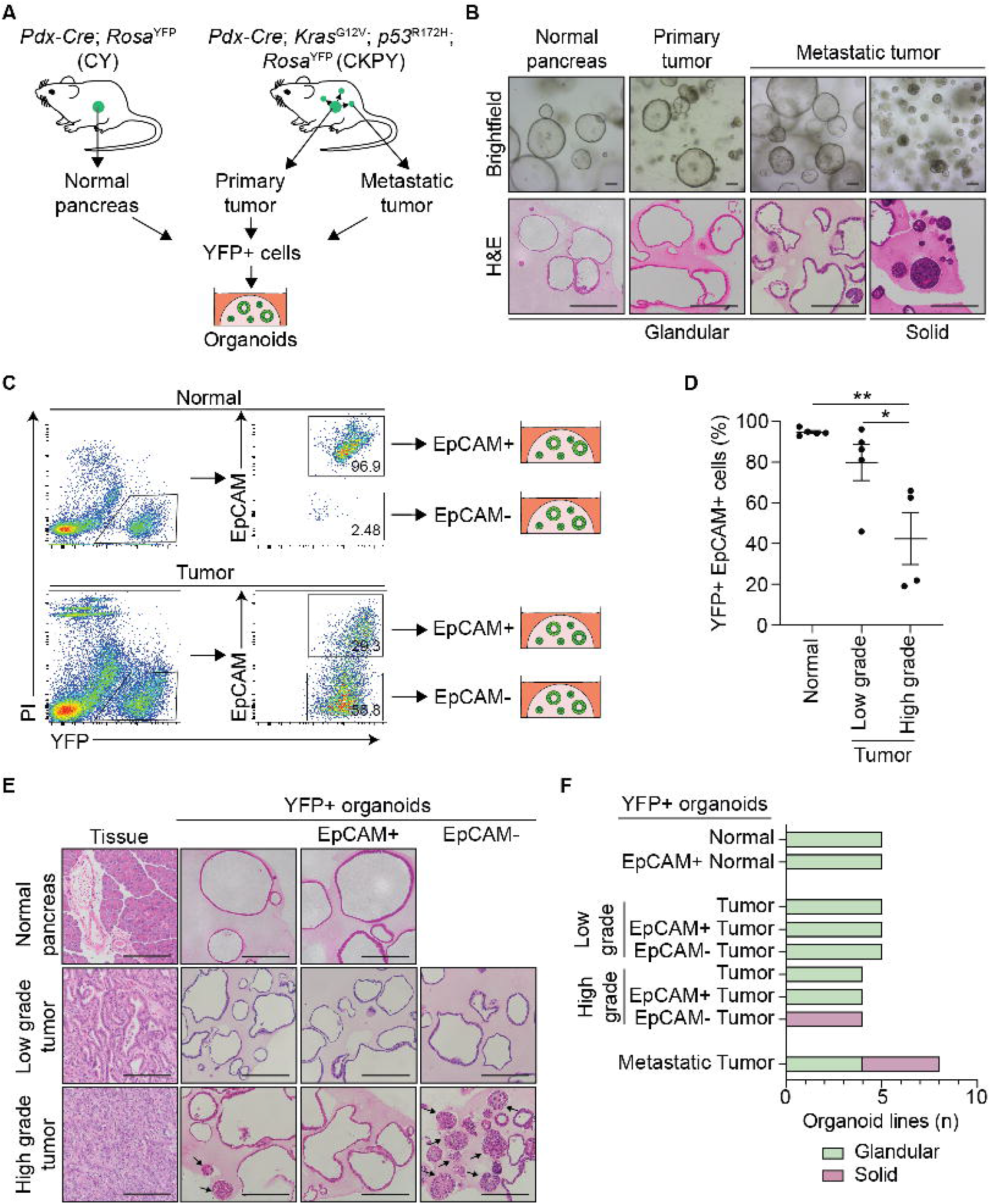
Generation of a murine organoid biobank with different morphological features. (A) Schematic representation of the workflow to generate organoids from primary and metastatic secondary tumors in the KPC model. YFP+ cells FACS isolated from *Pdx-Cre*; *Rosa*^YFP^ (CY) mice were used to generate normal pancreatic organoids, while YFP+ cells FACS isolated from either primary tumors or secondary tumors (liver, lung and diaphragm) from *Pdx-Cre*; *Kras*^G12V^; *p53*^*R*172H^; *Rosa*^YFP^ (CKPY) mice were used to generate tumor organoids. (B) Representative brightfield (top) and H&E (bottom) images of normal pancreatic, primary tumor and metastatic tumor organoids. Representative glandular and solid sub-type organoids are shown. Scale bar = 100 μm. (C) Schematic representation of the FACS gating strategy for isolation of YFP+EpCAM+ and YFP+EpCAM-tumor cells for the generation of organoids. (D) Percentage of live YFP+EpCAM+ cells isolated from the normal pancreas from CY mice (N=5), and either low grade tumors (N=5) or high grade (N=4) pancreatic tumors from CKPY mice. Each dot represents an individual mouse. Data is presented as mean +/- SEM. *p<0.05, Student’s t-test. (E) Representative H&E images of the primary tumor and corresponding YFP+ (left), YFP+EpCAM+ (middle) and YFP+EpCAM- (right) organoids. Scale bar = 100 μm. (F) Quantification of the organoid morphology for each MO line, represented as glandular (green) or solid (red). See also Figure S1, S2 and Table S1.

We observed two distinct MO morphological sub-types, denoted glandular and solid. H&E staining demonstrated that the glandular MO sub-type retained a single cellular layer with a central lumen, whereas the solid MO sub-type had multiple cellular layers often in the absence of a lumen (Figure 1B). There was no correlation between the tissue origin and morphological sub-types (Figure S2A). We questioned whether the differences in morphology that we observed were reflective of different states of EMT, following the observation that CKPY mice with high grade tumors often displayed regions with loss of expression of the epithelial marker EpCAM, while the same epithelial cells expressed the mesenchymal marker Vimentin consistent with a classic EMT phenotype (Figure S1B) (Aiello et al., 2017; Rhim *et al*., 2012b). Similar to other studies, we found that loss of E-Cadherin was not as consistent as the loss of EpCAM staining when examining EMT phenotypes (Simeonov et al., 2021)(Figure S1B).

In parallel to FACS sorting primary tumors for YFP+ cells, YFP+EpCAM+ and YFP+EpCAM-cells were also sorted to generate P-EMT MO sub-lines (Figure 1C). We found less YFP+EpCAM+ cells were present in the pancreas of mice retrospectively assigned as high-grade tumors by histopathology, consistent with the suggestion that some malignant cells have undergone EMT (Figure 1D). All of the YFP+EpCAM+ cells that were sorted from the normal pancreas of CY mice successfully formed MO; however, the low percentage of YFP+ EpCAM- cells isolated did not grow (Figures 1E and S2B). Within the established MOs, the YFP+EpCAM+ and the matched YFP+ normal pancreas MOs retained a glandular morphology across passages (Figure 1E-F, Figure S2C). All of the MOs established from the YFP+EpCAM+ and YFP+EpCAM-cells from the pancreatic tumors of CKPY mice successfully grew (Figures 1E-F and S2C). We noticed that the low grade primary tumors resulted in a higher percentage of glandular sub-type organoids in the YFP+ MOs; whereas the high grade tumors generated a mixture of solid and glandular sub-type MOs (Figures 1E-F and S2C). For the YFP+EpCAM+ and YFP+EpCAM-MO sublines, the low grade tumors generated glandular MOs (Figures 1E-F and S2C). Although, the YFP+EpCAM-MO from low grade tumors, were initially a solid sub-type they transitioned to a glandular sub-type after serial passaging (Figure S2C) suggesting they underwent MET in culture. In contrast, the YFP+EpCAM-MOs from high-grade tumors retained their solid morphology, suggesting differences in the plasticity in the MOs generated from either low grade or high grade tumors (Figure S2C).

### Murine organoid morphology is linked to elevated s100a4 and decreased s100a14 expression

In order to determine if the morphology of the organoids was linked to the EMT continuum we undertook a label-free based quantitative proteomics analysis of lysates generated from the YFP+EpCAM+ primary tumor, YFP+EPCAM-primary tumor and YFP+ secondary tumor MOs (Figure 2A, Table S2). Principle component analysis revealed a clear pattern of separation between the different MO sublines (Figure 2B). We next generated an EMT score based on a previously defined continuum of P-EMT signatures, namely: epithelial, EM-hybrid (H1-4), and mesenchymal (Simeonov *et al*., 2021). The glandular primary tumor YFP+EpCAM+ and glandular metastatic YFP+ MOs showed no clear association with an EM-hybrid or mesenchymal signature, while they were enriched for an epithelial signature (Figure 2C). The glandular YFP+EpCAM-MO generated from low grade primary tumors also had an enriched epithelial signature (Figure 2C). We found that the solid YFP+EpCAM- and solid YFP+ metastatic MOs had a lower epithelial signature, consistent with entry into a classic EMT state (Figure 2C). We also observed a lower H4 signature in the solid primary tumor YFP+EpCAM-organoids indicating P-EMT, while the solid YFP+ metastatic MOs had a higher mesenchymal signature, suggesting the metastatic organoids progressed towards C-EMT (Figure 2C).

**Figure 2.**
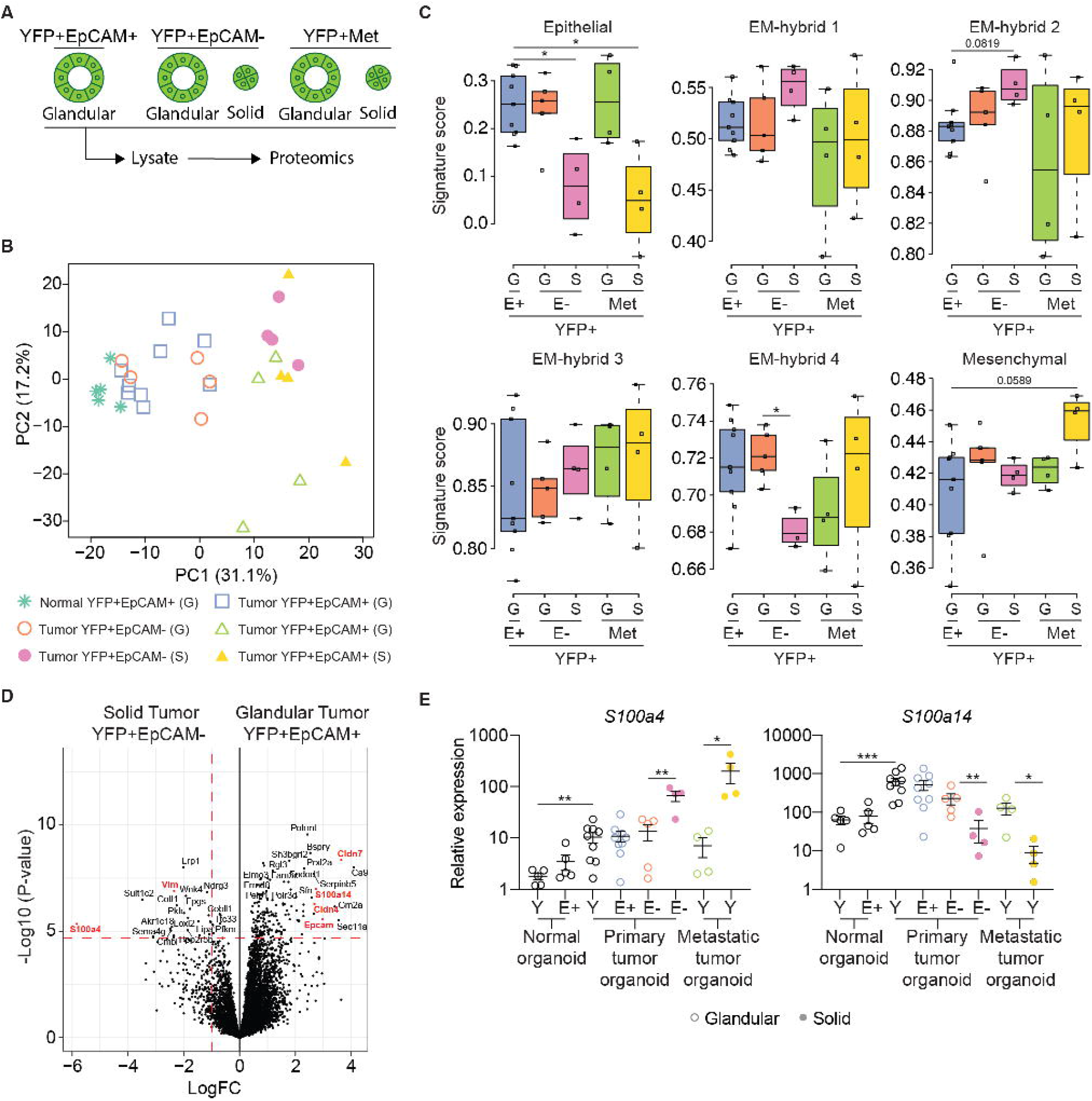
Murine organoid morphology correlates with P-EMT state. (A) Schematic representation of organoid lines used for label-free based quantitative proteomics analysis. Organoids with different morphologies were subjected to cell lysis followed by on-bead enzymatic digestion and subsequent mass spectrometry analysis. (B) Principal component analysis plot of 3,546 of the most variable proteins across organoid lines, including YFP+EpCAM+ normal (N=5), YFP+EpCAM+ tumor (N=9), YFP+EpCAM-tumor (N=9) and YFP+ secondary tumors (Met; N=8) organoids. The plot shows the separation of samples based on different principal components (PCs). Glandular (G) and solid (S) organoid lines are indicated. (C) The organoid signature score of the six previously described EMT subtypes (Simeonov *et al*., 2021) including YFP+EpCAM+ (E+; N=9), YFP+EpCAM- (E-; N=9) and YFP+ secondary (Met; N=8) tumor organoids. Glandular (G) and solid (S) organoid lines are indicated. (D) Volcano plot illustrating the log_2_ protein ratios in whole cell lysates of organoids, comparing glandular YFP+EpCAM+ with solid YFP+EpCAM-tumor organoids. Proteins were deemed differentially regulated in the log2 fold change in protein expression was > 1-fold and exhibited an adjusted *p*-value ≤ 0.05. (E) mRNA expression of *S100a4* and *S100a14* in YFP+ (Y), YFP+EpCAM+ (E+) and YFP+EpCAM- (E-) organoids. Each dot represents an individual organoid line. Glandular (open circle) and solid (closed circle) organoid lines are indicated. Data is relative to *Gapdh*, presented as mean +/- SEM. *p<0.05; **p<0.01; ***p<0.001, Mann-Whitney test. See also Figure S3.

We were curious as to whether our proteomics dataset could identify individual proteins that may be associated with morphological changes and EMT status. To this end, we first examined the differential expression between the glandular normal YFP+EpCAM+ and glandular tumor YFP+EpCAM+ MOs (Figure S3A) and found elevation of Agr2 in the tumor MOs, a protein in the TP53 pathway that has previously been shown to be expressed in human PDAC patients and linked to tumor cell dissemination (Dumartin et al., 2011). Other differentially expressed proteins included Nusap1, a tubulin protein involved in cell cycle regulation and Lgals4, a carbohydrate-binding protein (Figure S3A), both of which are associated with poor prognosis in PDAC patients (Deng et al., 2020; Hu et al., 2019). On examination of the differential protein expression between the glandular primary tumor MO P-EMT sub-lines, there were no major differences in protein expression (Figure S3B), while the glandular metastatic tumor MO had reduced Cldn7 compared to the glandular primary tumor MOs (Figure S3C).

We next compared the ‘epithelial’ glandular YFP+EpCAM+ and solid YFP+EpCAM-MO sub-lines, which revealed increased expression of EpCAM in the primary tumor glandular organoids, and increased Vimentin in the solid organoids, consistent with their P-EMT phenotypes (Figure 2D). In the solid organoids, there was also a significant reduction in the expression of Cldn7 and Cldn4, which belong to a tight junction protein family previously associated with tumor differentiation, liver metastasis and decreased survival in PDAC patients (Alikanoglu et al., 2015; Holczbauer et al., 2013; Soini et al., 2012). In contrast, the solid metastatic MOs had a reduction in Epcam and Cldn7 compared to the glandular primary tumor MOs (Figure S3D). Comparison of the solid and glandular YFP+EpCAM-sublines also revealed decreased expression of Cldn7 and Epcam in the solid MO (Figure S3E), while comparisons of the MOs from metastases did not reveal major differences from the solid MOs (Figure S3F-H). An overall comparison of the top 25 differentially expressed proteins (Figure 2D) within the MO sublines demonstrated that the expression profile among the MO was similar within morphology phenotypes (Figure S3I).

We were intrigued by the differential expression of S100a4 and S100a14 in our datasets (Figures S3I-K). S100 family proteins were recently found to be one of the most abundant secreted factors in PDAC when compared with the normal pancreas in both a murine model and patients (Tian et al., 2019). We observed differential expression of S100a4, S100a13, S100a14 and S100a16 between the glandular YFP+EpCAM+ and the solid YFP+EpCAM-organoids (Figure 2D, Figure S3J). Further examination of *S100a4* mRNA expression levels confirmed increased expression in YFP+ tumor MOs compared to YFP+ normal MOs, with a further significant increase in the solid MOs (Figure 2E). In contrast, while *S100a14* mRNA was significantly increased in YFP+ tumor MOs compared to YFP+ normal MOs, its expression was decreased in the solid MOs (Figure 2E). Publicly available transcriptomics data from pancreatic cancer patients and normal human pancratic tissue also confirmed the increase expression of both *S100A4* and *S100A14* in PDAC (Figure S4A) (Tang et al., 2017). We questioned whether the S100 proteins were associated with tumor grade and examined publicly available transcriptomics data from pancreatic cancer patients with high tumor purity (Raphael et al., 2017) and confirmed the association of *S100A4* with high grade tumors and *S100A14* expression with low grade tumors (Figure S3K).

### TGFβ1 induced solid morphology is associated with increased s100a4 expression

TGFβ1 has previously been associated with EMT (Gabitova-Cornell *et al*., 2020; Su *et al*., 2020). In patients, a TGFβ1 signature is significantly elevated in high grade tumors (Figure S4B). In the CKPY mice we also observed that *tgfb1*, along with its receptor *tgfbr2*, was increased in the high grade murine tumors (Figure S4C). However, within the P-EMT MO sub-lines, we did not observe a significant increase in the expression of endogenous *tgfb1*, rather a trend in decreased expression along with a significant reduction in *tgfbr2* expression in the solid MOs (Figure S4D).

Since a contribution of TGFβ1 to EMT may be contextual, we mimicked the high levels of TGFβ1 secreted by the tumor microenvironment (Bernard et al., 2019; Elyada et al., 2019) *in vitro* through treatment of MO cultures with recombinant TGFβ1 (Figure 3A). We observed that the glandular MOs adopted a solid morphology following TGFβ1 treatment, while a portion of the solid organoids began to display an invasive ‘branching’ phenotype (Figure 3B-C). These TGFβ1 induced morphological changes were associated with a significant increase in the expression of classic mesenchymal markers, including *Zeb1, Snai1/2, Vim, Cdh2 and Fn1* (Figure 3D) suggesting the induction towards a P-EMT phenotype. We also observed a significant increase in the expression of *S100a4* and a significant decrease in the expression of *S100a14* (Figure 3E) placing regulation of the expression of these genes these downstream of the TGFβ1 pathway.

**Figure 3.**
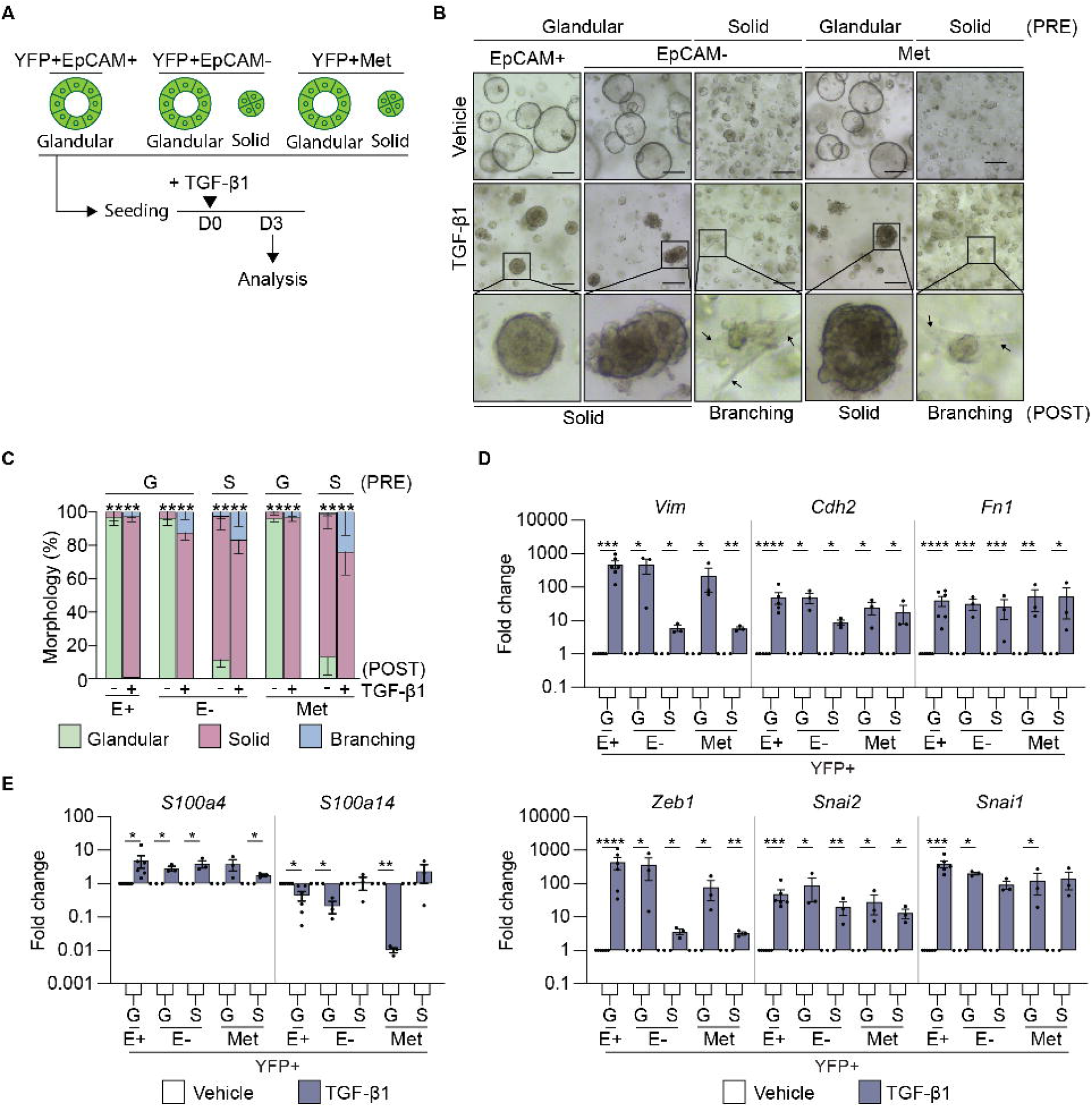
TGF-β1 treatment induces a mesenchymal signature in tumor organoids. (A) Schematic representation of the treatment of murine tumor organoids with recombinant TGFβ1 and timing of quantification of the morphology and gene expression signatures. (B) Representative brightfield images of YFP+EpCAM+ (N=6), YFP+EpCAM- (N=6) and YFP+ secondary (Met, N=6) tumor organoids on day 3 post TGFβ1 treatment. The pre-treatment (PRE) organoid morphology is indicated. Representative images of organoids post-treatment (POST) are shown. Arrows indicate a branching phenotype. Images are representative of 3 biological replicates. Scale bar = 300 μm. (C) Quantification of the tumor organoid morphology 3 days post TGFβ1 treatment. YFP+EpCAM+ (N=6), YFP+EpCAM- (N=6) and YFP+ secondary (Met; N=6) tumor organoids. Organoids are grouped as glandular (G) and solid (S) morphologies pre-treatment (PRE). Quantification of glandular (green), solid (red) and branching (blue) post cytokine treatment (POST). Data includes 3 biological replicates and is presented +/- SEM. ****p<0.0001, Chi-square test. (D-E) mRNA expression levels of mesenchymal markers, *Vim, Cdh2, Fn1, Snai1, Snai2, Zeb1* (D); and S100 family members, S*100a4, S100a14* (E), in YFP+EpCAM+ (E+), YFP+EpCAM- (E-) primary tumor organoids and YFP+ secondary tumor organoids following the addition of TGFβ1 (blue). Each dot represents an individual organoid line. Organoids are grouped as glandular (G) and solid (S) morphologies. Data includes 3 biological replicates and is presented as log10 fold change relative to the vehicle (white) control, mean +/- SEM. *p<0.05, **p<0.01, ***p<0.001, ****p<0.0001, paired t-test. See also Figure S4.

Given the adverse outcomes observed following direct targeting of TGFβ1 in pancreatic cancer patients (Alvarez et al., 2019), we explored whether other cytokines downstream of TGFβ1, that may be more favorable therapeutic targets, were implicated in the EMT-associated changes in morphology that we observed. To this end, we confirmed that following treatment of MO cultures with recombinant TGFβ1 (Figure 4A), the expression of the IL-6 family cytokines *il6, lif* and *il11* were significantly increased (Figure 4B). Interestingly, each of these cytokines, or their receptors, have been implicated in EMT processes (Lee et al., 2019; Shi et al., 2019). In publicly available transcriptomics datasets, *IL6* was associated with high grade tumors in patients and *LIF* was associated with low grade tumors (Figure S5A). We hypothesized that autocrine IL-6 family cytokine signaling would be augmented in TGFβ1 treated MOs, as we also observed increased expression of the shared signaling receptor *il6st* (gp130; Figure 4C). We were curious as to whether a hierarchy of importance may exist within this cytokine family, as we observed increased expression of the cognate cytokine receptor, *il6ra* and decreased expression of *lifr* in the TGFβ1 treated MOs (Figure 4D). We did not observe expression of *il11ra* in the tumor MOs (Figure 4D), contrary to previous published reports (Ohlund et al., 2017). Consistent with the activation of an autocrine IL-6 signaling cascade, *Socs3*, a STAT3 target gene that acts as a negative physiological regulator of signalling, was also increased following stimulation with recombinant TGFβ1 (Figure 4E).

**Figure 4.**
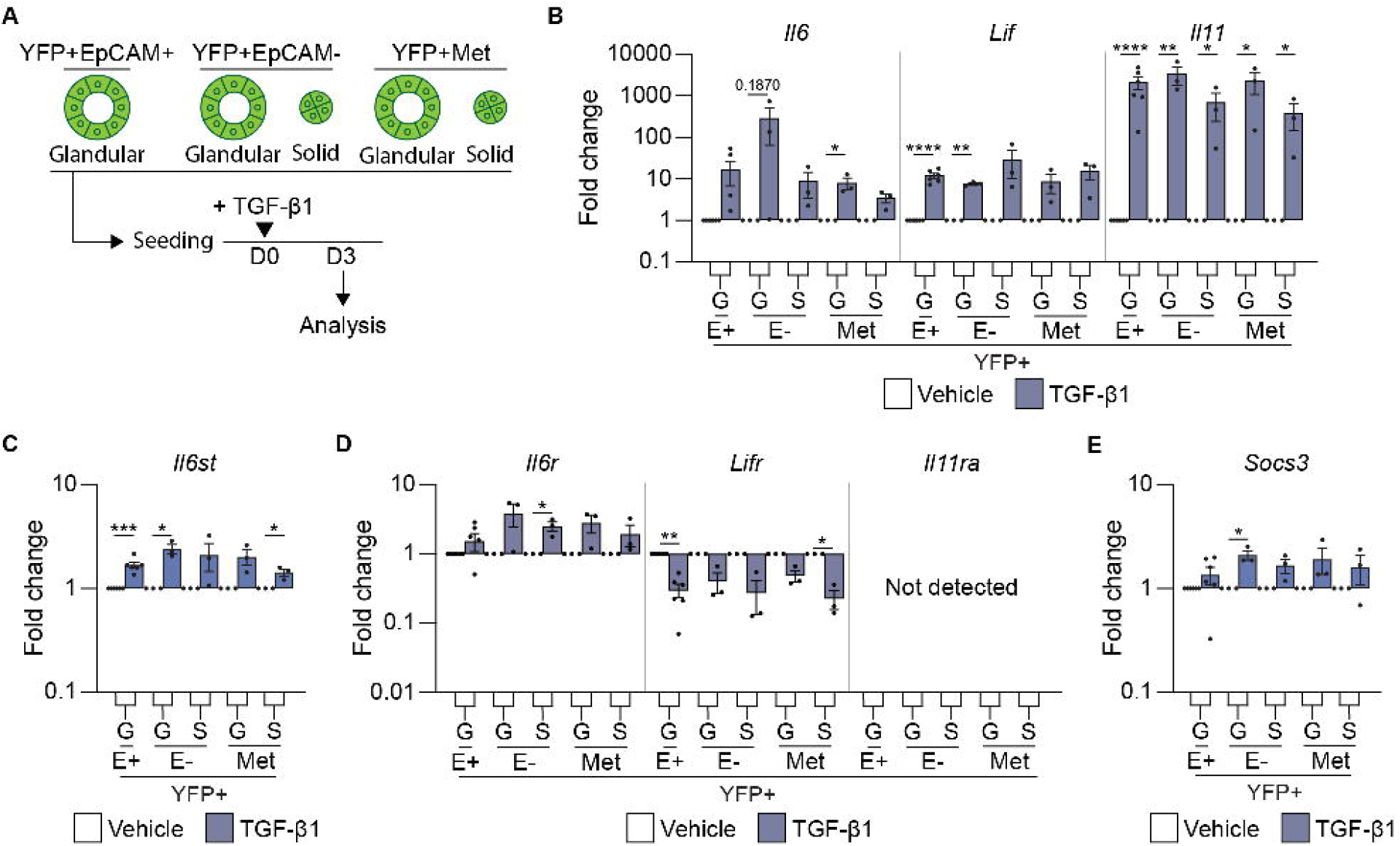
TGFβ1 induces the expression of IL-6 family cytokines. (A) Schematic representation of the treatment of murine tumor organoids with recombinant TGFβ1 and timing of quantification of gene expression signatures. (B-E) mRNA expression levels of IL-6 family cytokines, *Il6, Lif*, and *Il11* (B); IL-6 family cytokine receptors *il6st* (C); *Il6r, Lifr*, and *Il11r* (D), and the IL-6 cytokine family target gene, *Socs3* (E), 3 days post treatment with TGFβ1. Each dot represents an individual organoid line. Organoids are grouped as glandular (G) and solid (S) morphologies. Data includes 3 biological replicates and is presented as log10 fold change relative to the vehicle (white) control, mean +/- SEM. *p<0.05, **p<0.01, ***p<0.001, ****p<0.0001, paired t-test.

In order to determine if IL-6 and LIF directly induced EMT processes, we next cultured the MOs in the presence of either recombinant IL-6 or recombinant LIF (Figure 5A); however, we did not observe a change in MO morphology (Figure 5B-C). Consistent with a lack of EMT induction, we did not observe an increase in the expression of classic mesenchymal markers (Figure S5B) or *s100a4*, rather we observed an increase in *s100a14* (Figure 5D), which has previously been associated with IL-11 signaling (Al-Ismaeel et al., 2019). We did observe an increase in the STAT3 target gene, *Socs3*, highlighting successful activation of the signaling pathway (Figure S5C). These results indicate that the TGFβ1 mediated EMT process that is associated with a solid MO morphology and elevated *s100a4*, is not mediated by members of the IL-6 cytokine family.

**Figure 5.**
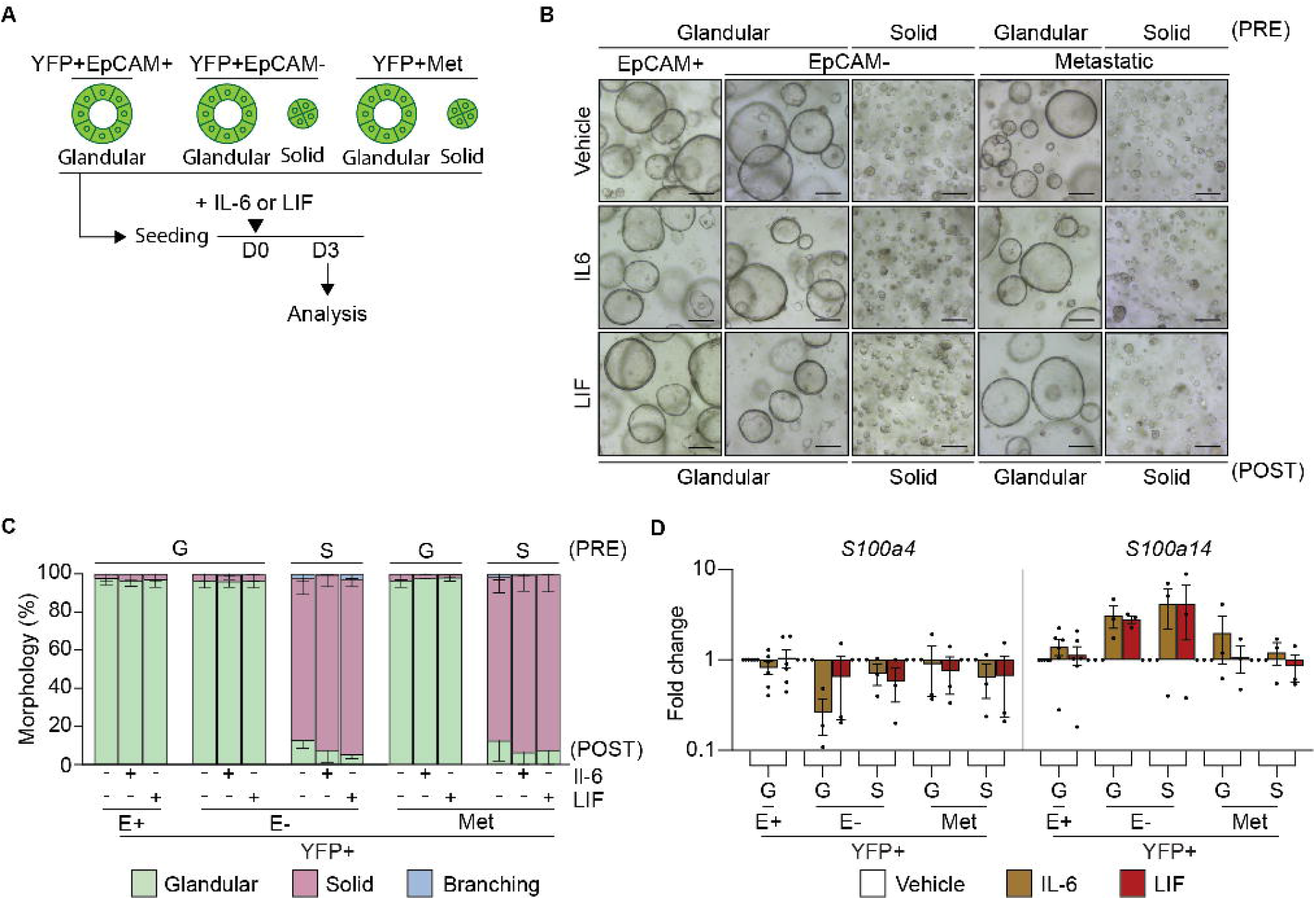
IL-6 or LIF do not alter tumor organoid morphology or induce EMT. (A) Schematic representation of the treatment of murine organoids with recombinant IL-6 or LIF and timing of quantification of the morphology and gene expression signatures. (B) Representative brightfield images of YFP+EpCAM+ (E+; N=6), YFP+EpCAM- (E-) and YFP+ secondary (Met; N=12) tumor organoids on day 3 post treatment (POST) with the indicated cytokine. Organoids are grouped as glandular (G) and solid (S) morphologies pre-treatment (PRE). Scale bar = 300 μm. (C) Quantification of the tumor organoid morphology 3 days post treatment with the indicated cytokine. YFP+EpCAM+ (E+, N=6), YFP+EpCAM- (E-, N=12 primary and secondary). Glandular (green), solid (red) and branching (blue). Data is presented +/- SEM. Results are not significant, Chi-square test. (D) mRNA expression levels of S100 family proteins, S*100a4* and *S100a14*, in YFP+EpCAM+ (E+), YFP+EpCAM- (E-) primary tumor organoids and YFP+ secondary tumor organoids following the addition of the indicated cytokine (IL-6, brown; LIF, red). Each dot represents an individual organoid. Organoids are grouped as glandular (G) and solid (S) morphologies. Data includes 3 biological replicates and is presented as log10 fold change relative to the vehicle (white) control, mean +/- SEM. paired t-test. See also Figure S5.

### Solid organoids form high grade allograft tumors with increased s100a4 expression

Since we observed in our own patient cohorts that high levels of serum TGFβ1 in PDAC patients increased with the stage of the tumour (Figure S6A) we further explored the relationship between TGFβ1, organoid morphology and high grade tumors by establishing MO allograft models. To this end, YFP+EpCAM+ primary tumor, YFP+EpCAM-primary tumor and YFP+ secondary tumor MOs were engrafted subcutaneously into C57B/l6 mice (Figure 6A). We found that 100% of the MOs with a solid morphology engrafted, while the glandular MOs had a variable engraftment success rate (Figure S6B).

**Figure 6.**
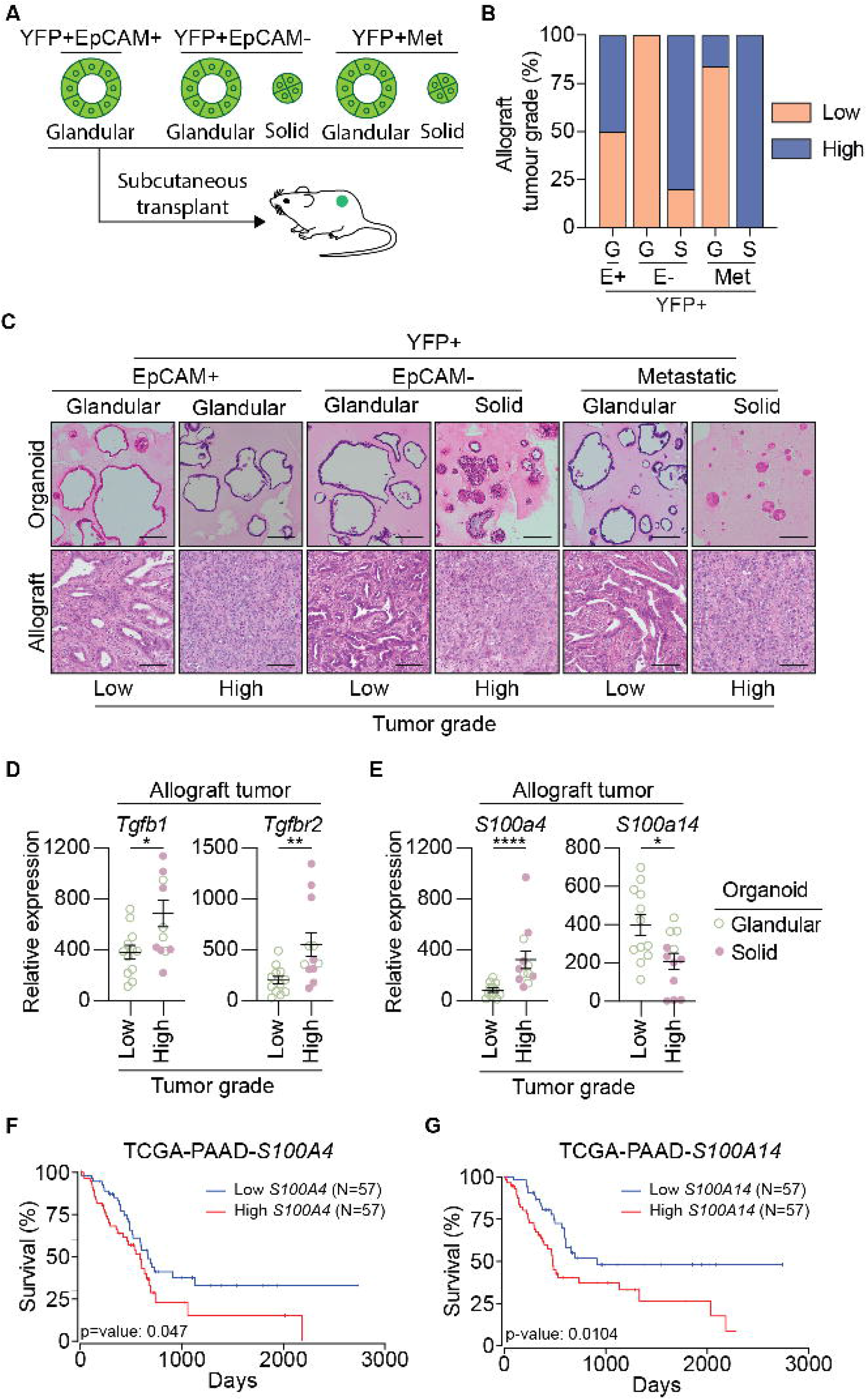
Organoid-derived allografts resemble original tumors and exhibit an EMT phenotype. (A) Schematic representation of the subcutaneous transplant of organoids into gender matched wild-type C57BL/6 mice to generate allograft lines. (B) Quantification of the number of low grade (orange) and high grade (blue) allograft tumors, generated from YFP+EpCAM+ (E+), YFP+EpCAM- (E-) primary tumor organoids and YFP+ secondary tumor organoids. Organoids are grouped as glandular (G) and solid (S) morphologies. (C) Representative H&E images of the tumor organoid, and corresponding allograft generated from YFP+EpCAM+ (E+), YFP+EpCAM- (E-) primary tumor organoids and YFP+ secondary tumor organoids. Organoids are grouped as glandular (G) and solid (S) morphologies. (D-E) mRNA expression of *Tgfb1, Tgfbr2* (D), and the S100 family proteins, S*100a4, S100a14* (E), in low grade (N=4 tumors) and high grade (N=4 tumors) allograft tumors derived from YFP+EpCAM+ MO, YFP+EpCAM-MO, and YFP+ secondary tumor MO. Each dot represents an individual tumor. Data is relative to *Gapdh*, presented as mean +/- SEM. *p<0.05, **p<0.01, Mann-Whitney test. (F) Kaplan-Meier plots of overall survival for PDAC patients based on high (top 33%) and low mRNA expression (bottom 33%) of *S100A4* and *S100A14* obtained from the TCGA-PAAD dataset. Plots and p-value were determined using the OncoLnc web tool (Anaya, 2016). See also Figure S6.

Retrospective histopathological analysis of the resulting allograft tumors indicated that nearly all of the solid organoids formed high grade tumors, while a mixture of low and high grade tumors were derived from the glandular MOs (Figure 6B-C). The high grade allograft tumors had classic features of EMT, including loss of EpCAM expression and increased Vimentin expression (Figure S6C). We did observe that some of the glandular YFP+EpCAM-MO re-expressed EpCAM in the allograft tumours, suggesting MET occurred following transplantation of those MO sub-lines (Figure S6C). We observed higher expression of both the TGFβ1 cytokine and its cognate receptor in the high grade allograft tumors (Figure 6D). These observations corresponded with increased expression of *S100a4* in the high-grade allograft tumors, while *S100a14* expression was decreased (Figure 6E).

Taken together, we have revealed that MOs with a solid morphology have an P-EMT phenotype, which can be augmented by TGFβ1 and transitioned to a C-EMT phenotype. We also show that solid organoids form allografts that are high grade tumors, with high expression of both TGFβ1 and *s100a4*. Our findings link TGFβ1 with changes in the expression of *S100a4* and *S100a14*, which individually are associated with poor survival in PDAC patients (Figures 6F-G). Our results suggest that S100A4 may represent an important biomarker to predict EMT state, disease progression and survival.

## DISCUSSION

The convergence of tumor intrinsic and microenvironment signals that promote the transition through cancer cell EMT states, and the reversal of these processes, will inevitably impact tumor progression (Nieto et al., 2016). Given the highly aggressive nature of PDAC, the identification of biomarkers that could predict advanced disease would be clinically useful.

We have employed a murine model of PDAC enabled by oncogenic recombination in epithelial cells (Rhim *et al*., 2012b). We observed that more than half of the primary tumors that arose displayed EMT features characterized by the gain of expression of mesenchymal markers, and the loss of expression of the epithelial marker, EpCAM. Our study builds on the recent technical advances realized through the generation of a murine organoid biobank from primary and secondary tumors (Boj et al., 2015; Messal et al., 2019; Roe et al., 2017; Wang *et al*., 2019), by including a YFP reporter allele that allowed us to isolate pure epithelial cells and generate matched MOs from secondary tumors present in multiple organs, including the lung (Roe *et al*., 2017). Moreover, the addition of FACS gating strategies using EpCAM as a reliable EMT lineage marker (Gires et al., 2020; Liao and Yang, 2020; Pastushenko et al., 2021; Simeonov *et al*., 2021) permitted the generation of P-EMT sub-lines providing an opportunity to better understand the spectrum of P-EMT programs, which has previously been technically challenging.

The emergence of different morphologies within our MO biobank permitted phenotyping of cell intrinsic behaviors in a way not possible with monolayer cell lines (Aiello *et al*., 2018). Similar to previous reports for patient-derived PDAC organoids (Driehuis et al., 2019; Huang *et al*., 2020a; Romero-Calvo et al., 2018; Seino et al., 2018; Sharick et al., 2020; Tiriac et al., 2018), we showed that solid murine organoids were often derived from high grade tumors. Our proteomics analysis revealed that the morphology of a PDAC organoid correlated with the continuum of EMT states, which has previously been reported for mammary MOs (Jung et al., 2019; Shamir et al., 2014). We found that within the MOs isolated from both primary and secondary tumors, glandular MOs were often in an epithelial state, while solid MOs had often undergone P-EMT. We also found that the P-EMT MOs gave rise to different EMT spectrums along a previously described trajectory of epithelial, EMT-hybrid and mesenchymal states (Simeonov *et al*., 2021). Interestingly, following serial passaging and allograft generation a sub-set of the solid organoids with a P-EMT phenotype underwent MET, characterized by a reversion to a glandular phenotype and the re-expression of EpCAM. This phenomenon is not uncommon, and has previously been reported *ex vivo*, following the culture of P-EMT cell lines (Aiello *et al*., 2018) and *in vivo* following the migration of cells from primary tumors to distant organs (Beerling *et al*., 2016; Reichert *et al*., 2018).

Among the proteins identified within a P-EMT state in solid MOs was the S100 protein family, which consists of 25 members that have a variety of intracellular and extracellular cellular functions including calcium homeostasis, proliferation and apoptosis (Wang et al., 2018). S100 proteins have also been shown to interact with cytoskeletal proteins thus affecting cellular morphology and migration (Gires *et al*., 2020; Wang *et al*., 2018). We were intrigued by the S100 proteins, as they have previously been shown to be secreted by epithelial cells in response to tissue damage or inflammatory responses (Tian *et al*., 2019; Xia et al., 2018). We found that S100a14 was increased in the MOs derived from PDAC tumors compared to the normal MOs, which is consistent with the increased expression observed in murine cancer cells, patient cancer cell lines, and link to poor overall survival (Al-Ismaeel *et al*., 2019; Hosein et al., 2019; Simeonov *et al*., 2021). S100a14 has been described as a mesenchymal marker, since upregulation of the transcription factor Gli1 promotes EMT and increased s100a14 expression (Xu et al., 2014). However, we and others have shown using murine models (Hosein *et al*., 2019) and human cell lines (Al-Ismaeel et al., 2019) that that s100a14 is associated with epithelial and not EMT states, since its expression was lower in glandular MOs, murine and patient low grade tumors.

We were also interested in the increased expression of s100a4 that we observed was associated with P-EMT in solid organoids, since s100a4 has also been associated with poor survival in PDAC (Dreyer et al., 2020). S100a4 has also been shown to play a role in EMT, metastasis, and is highly expressed in murine PDAC mesenchymal tumor cells, patient cell lines, and PDAC patient high grade tumors suggesting a clear association with EMT (Al-Ismaeel *et al*., 2019; Chen et al., 2015; Dreyer *et al*., 2020; Simeonov *et al*., 2021; Xu *et al*., 2014). As s100a14 is also increased in tumor cells, and associated with poor prognosis, s100a4 may be a more useful biomarker of advanced disease. Interestingly, high *S100A4* expression correlates with resistance to Gemcitabine, the mainstay chemotherapy provided to PDAC patients (Ma et al., 2015; Mahon et al., 2007), with EMT also known to alter response to chemotherapy (Creighton et al., 2009; Shibue and Weinberg, 2017).

Secreted factors within the tumor microenvironment are known to influence EMT processes in PDAC (Ligorio et al., 2019). We explored the contribution of TGFβ1, and found that recombinant TGFβ1 could transition a glandular organoid to a solid phenotype, which was associated with the induction of a classic EMT gene signature. As previously shown in PDAC patient derived organoids (Huang *et al*., 2020a), we observed that a branching phenotype could also be induced by recombinant TGFβ1 suggesting a transition to a C-EMT state. We were particularly intrigued by the observation that TGFβ1 could upregulate *S100a4* consistent with other studies (Chen *et al*., 2018; Krebs *et al*., 2017). It is thought that TGFβ1 may regulate S100a4 through the EMT-TF, Zeb1, as Zeb1 overexpression also resulted in an increase in *S100A4* expression in cell lines, while *S100A14* decreased consistent with our observations (Al-Ismaeel et al., 2019).

Despite being an obvious therapeutic target to prevent transition from early stage EMT to a C-EMT, there is likely limited clinical benefit to be had following targeting of TGFβ1 in cancer due to the high probability of on-target toxicities due to the complexities of TGF signalling and function within the immune system. For this reason, we explored the role of IL-6 family cytokines in EMT, which are readily targeted therapeutically, and which we and others have shown are induced downstream of TGFβ1 signalling (David et al., 2016; Su et al., 2020). However, we failed to detect changes in MO morphology following stimulation with recombinant IL-6 family cytokines, nor did we detect an increase in classic EMT markers suggesting that this cytokine family alone does not promote EMT. Moreover, despite previous reports that IL-11 can upregulate *S100A4* and *S100A14* in human PDAC cell lines (Al-Ismaeel et al., 2019) we failed to detect IL-11Rα_1_ expression in our MO biobank. Other studies have also shown that treatment of lung cancer cell lines with IL-6 alone did not induce EMT, although the presence of receptor components was not confirmed (Liu et al., 2014). We also did not observe an induction of *S100a4* following stimulation with IL-6 or LIF, but rather we observed a significant induction of *S100a14* suggesting that while IL-6 and LIF may contribute to tumor features they do not directly contribute to EMT. Thus, therapeutic inhibition of the IL-6 cytokine family alone is unlikely to prevent EMT. However, activation of the signal transducer and activator of transcription (STAT)-3, a pro-tumorigenic transcription factor downstream of the IL-6 family of cytokines, significantly correlated with *S100A4* expression in PDAC patient tissue (Al-Ismaeel et al., 2019) suggesting cooperative induction of *S100A4* is possible. Previous reports have also suggested synergy between the ZEB1 EMT-TF and the IL-6 family cytokines results in augmentation of *S100A4* expression, which was not explored here (Al-Ismaeel et al., 2019).

Taken together, our observations suggest that both extracellular and cell intrinsic C-EMT programs converge on S100A4, which may represent a useful biomarker to predict disease progression and prognosis.

## AUTHOR CONTRIBUTIONS

RRJL and TLP conceived the study, designed the experiments, and wrote the manuscript. RRJL performed all MO experiments. KYF, AP contributed to *in vivo* experiments. HG analysed the public transcriptomic datasets and proteomics dataset. JY, LFD and RL completed the proteomics analysis. BL and KYF performed serum analysis and PG provided human samples. NK, AWB and MDWG contributed critical reagents and intellectual input. FH, MDWG and SMG supervised experiments and provided intellectual input. All authors have read and agreed to the manuscript.

## ACKNOWLEDGEMENTS

We wish to thank the WEHI bioservices and histology facilities for excellent technical support. This work was supported by generous donations from the Philip Hemstritch Pancreatic Cancer Research Program and Donald Cant Watts Corke, Australia. TLP is a Sylvia and Charles Viertel Charitable Foundation Senior Medical Research Fellow. Funding from the Victorian State Government Operational Infrastructure Support Scheme is acknowledged.

## RESOURCE AVAILABILITY

### Lead contact

Further information and requests for resources and reagents should be directed to and will be fulfilled by the lead contact, Tracy Putoczki.

### Materials availability

All unique materials generated in this study are available from the lead contact with a completed materials transfer agreement.

### Data and code availability

The mass spectrometry proteomics data have been deposited to the ProteomeXchange Consortium via the PRIDE partner repository with the dataset identifier PXD030992.

## EXPERIMENTAL MODEL AND SUBJECT DETAILS

### Mouse Strains

The *Pdx*^Cre^; *Rosa*^YFP^ (CY) and *Pdx*^Cre^; *Kras*^G12V^; *p53*^R172H^; *Rosa*^YFP^ (CKPY) mice (Rhim et al., 2012a) were maintained on a C57BL/6 background and bred and maintained in a specific pathogen free animal facility at WEHI. All experiments involving mice were approved by the WEHI Animal Ethics Committee (AEC approval #2019.015 and #2020.032). CKPY mice were aged between 12 - 27 weeks and collected together with aged and gender matched CY mice.

### Generation of Organoids

FACs sorted cells from murine tissue were resuspended in >90% v/v Matrigel (Corning) and grown in murine pancreatic organoid medium (MPOM) containing advanced DMEM/F12 (Gibco) containing 10 mmol/L HEPES (Gibco), 1X GlutaMAX (Gibco), 1x penicillin/streptomycin (Gibco), 5% v/v Rspo2-Fc conditioned medium (harvested from transiently transfected FreestyleTM-293F cells, Thermofisher), 5% v/v Noggin conditioned medium (harvested from Noggin-expressing 239 cells obtained from Foundation Hubrecht Organoid Technology (HUB), Hubrecht Institute, Uterecht, The Netherlands), 10 mmol/L nicotinamide, 1% v/v B-27 supplement without Vitamin A, 1 mmol/L N-acetyl-L-cysteine, 100 ng/mL rh FGF-10, 50 ng/mL rh EGF, 10nmol/L rh [Leu15]-gastrin I, 3 μmol/L prostaglandin E2. Following passaging, the organoids were cultured in MPOM with 10 μM Y-27632 (Sigma) and 5 μM GSK-3 inhibitor (Sigma) for the first 3 days, followed by culturing in normal MPOM for regular maintenance as described previously (Broutier et al., 2016).

### Generation of Allografts

Wild-type C57BL/6 mice were used to establish murine organoid-derived allografts. Wild-type C57BL/6 mice were bred and maintained in a specific pathogen free animal facility at WEHI. All experiments involving mice were approved and monitored by the WEHI Animal Ethics Committee (AEC approval #2017.033 and ##2020.032).

To generate allografts, one confluent well of organoids (approximately 50,000 cells) was resuspended in 100 μL of 50% v/v PBS/ 50% v/v Matrigel and subcutaneously injected into each flank of C57BL/6 mice (gender matched to the organoid). The tumor length (l) and width (w) were measured weekly using digital calipers and tumour volumes were calculated using the formula: 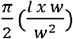. When the combined volume of both flanks reached 1 cm^3^, the mice were euthanised, and tissues were collected. If no visible tumour was present 6 months post engraftment, the engraftment was deemed unsuccessful and the mouse was euthanised.

### Patient Serum

Blood was collected from de-identified healthy or PDAC patients that had consented to a project governed by WEHI (G16/05) and serum collected using a Sarstedt tube, at 400 g for 15 min for storage at -80 °C.

## METHOD DETAILS

### Fluorescence-Activated Cell Sorting

Pancreas, liver and lung were digested in media containing 240units/mL Nystatin, 5μg/mL Amphotericin B, 1% w/v FBS, 0.125 mg/mL Collagenase, 0.125 mg/mL Dispase II, 0.1 mg/mL DNAse I, 1x advanced DMEM/F12, 1x penicillin/streptomycin, 1x GlutaMAX, 10mM HEPES for 30 min at 37 °C. Digested samples were passed through a 70μm strainer and washed with FACS buffer (PBS with 3% v/v FBS and 1% v/v EDTA). Pancreas, liver, lung and blood samples were incubated in red blood cell lysis buffer (ThermoFisher Scientific), washed and stained with FITC-fluorochrome conjugated Epcam antibody (Abcam, 1:1000) at 4 °C for 20 min. Propidium iodide (PI, ThermoFisher Scientific; 1:10,000) was added to the samples prior to FACS. Following preparation of the sample, FACSAria (BD Biosciences) sorting was performed.

### Histology

Murine tissue were fixed in 10% v/v formalin overnight, while organoids were fixed in 10% v/v formalin containing 0.5% v/v Glutaraldehyde (Sigma) for 45 min, embedded in paraffin and sectioned at a thickness of 4 μm. Staining of haematoxylin and eosin (H&E) was performed using standard procedures.

### Immunohistochemistry

Unstained paraffin sections were dewaxed, rehydrated and heat induced antigen retrieval performed by incubating the slides in Tris-EDTA-Tween 20 buffer (10 mM Tris base, 1 mM EDTA, 0.05% v/v Tween 20, pH 9.0, pH 9) or citrate buffer (10 mM, pH 6, ThermoFisher Scientific) as required for each primary antibody. To block endogenous peroxidases, slides were incubated in 3% v/v hydrogen peroxide (biolab) for 20 min followed by washes in ddH2O and TBST. Sections were incubated with blocking buffer (TBST with 5% v/v normal goat serum), in a humidified chamber for 1 hr at RT. After blocking, the sections were incubated in the desired primary antibody diluted in blocking buffer in a humidified chamber overnight at 4°C. The slides were then washed and incubated in the appropriate secondary antibody diluted in blocking buffer for 1 hr at RT.

To visualise staining, the slides were incubated in 3, 3’-Diaminobenzidine (DAB, Agilent) and then counterstained with haematoxylin (Abcam), dehydrated and mounted with Histomount (Life technologies).

### Imaging

Brightfield images of the organoids, histology and histochemical slides were taken on an Olympus CX23 upright light microscope (Olympus), with an Olympus DP22 camera, using CellSens Entry (Olympus) software. Organoid morphology quantification of either glandular of solid were manually quantified using ImageJ.

### RNA isolation, cDNA synthesis and qPCR

Tissue samples were homogenized in TRIzol (Ambion) using a Tissue Lyser II (Qiagen). 1 mL of chloroform (Merck Milipore) was added into each tube, mixed gently and centrifuged for 15 min at 3,000 rpm at 4°C. The upper layer, was transferred into a 1.5 mL tube (Eppendorf) containing 350 μL of 100% v/v isopropanol. RNA extraction was performed using a Qiagen RNA minikit (Qiagen) according to manufacturer’s protocol. PDOs were grown until confluence and released from matrigel by incubating in 500 μL of organoid harvesting solution (Gibco) at 4 °C with gentle rocking for 45 min. The samples were washed and RNA extraction was performed using Qiagen RNA minikit (Qiagen) according to manufacturer’s protocol. cDNA was generated using the high-fidelity cDNA synthesis kit (Applied Biosystem) as per manufacturer instructions.

The qRT-PCR was performed on a Viia 7 real-time PCR system with the following steps: initial denaturation for 10 min at 95 °C, denaturation for 20 s at 94 °C, annealing for 15 s at 60 °C and extension for 15 s at 72 °C for 40 cycles using gene-specific Taqman probes (Applied Biosystems). A relative comparative threshold (CT) method was used in the analysis of qRT-PCR using Microsoft Excel (Microsoft). The ΔCT values were calculated by subtracting the CT of the gene of interest from the CT of the housekeeping genes. The ΔCT was then calculated to determine the individual normalised values (2-ΔCT). Results for organoids were presented as fold change to matched YFP+ organoids. Results for cytokine stimulation were presented as fold change to vehicle.

### Cytokine stimulations

Organoids were plated in 2 wells in a 24-well plate for each condition. Vehicle (PBS) and recombinant cytokines including 10 ng/mL of TGFβ1, 100 ng/mL IL-6 or 100 ng/mL LIF were diluted in MPOM and added to the culture on day 0. On day 3 post stimulation, brightfield images of the organoids were taken for quantification and harvested for qPCR.

### Mass spectrometry-based proteomics

Organoids were lysed in RIPA buffer containing 100 mM NaCl, 10 mM Tris-HCl, 1% (v/v) glycerol, 50 mM NaF, 2 mM EDTA, 1% (v/v) Triton X-100, 1 mM Na3VO4, Complete mini protease inhibitor cocktail and Complete mini phosphatase inhibitor and 20 μg per replicate were prepared for proteomics analysis using the USP3 protocol previously described (Louis et al., 2020) with some minor modifications. Lysates were heated at 95°C for 10 min in a buffer containing 1% (v/v) SDS, 100 mM Tris (pH 8), 10 mM TCEP (Tris (2-carboxyethyl) phosphine) and 40 mM 2-chloracetamide and. Magnetic PureCube Carboxy agarose beads (Cube Biotech) were added to all samples along with acetonitrile (70% v/v final concentration) and incubated at room temperature for 20 mins. Samples were placed on a 96-well magnetic rack, supernatants discarded, and the beads washed twice with 70% ethanol and once with neat ACN. ACN was completely evaporated from the tubes using a CentriVap (Labconco) before the addition of digestion buffer (50 mm Tris) containing 0.8 μg Lys-C (Wako, 129–02541) and 0.8 μg Trypsin-gold (Promega, V5280). Enzymatic digestions proceeded under agitation (400 rpm) for 1 h at 37°C. Following digestion, samples were transferred to preequilibrated C18 StageTips for sample clean-up. The eluates were lyophilized to dryness using a CentriVap (Labconco), before reconstituting in 60 μl 0.1% FA/2% ACN ready for mass spectrometry analysis.

Peptides (2 μl) were separated by reverse-phase chromatography on a C_18_ fused silica column packed into an emitter tip (IonOpticks), using a nano-flow HPLC (M-class, Waters). The HPLC was coupled to a timsTOF Pro (Bruker) equipped with a CaptiveSpray source. Peptides were loaded directly onto the column at a constant flow rate of 400 nL/min with buffer A (99.9% v/v Milli-Q water, 0.1% v/v FA) and eluted with a 90-min linear gradient from 2 to 34% buffer B (99.9% v/v ACN, 0.1% v/v FA). The timsTOF Pro was operated in PASEF mode using Compass Hystar 5.1. Settings were as follows: Mass Range 100 to 1700m/z, 1/K0 Start 0.6 V·s/cm^2^ End 1.6 V·s/cm^2^, Ramp time 110.1ms, Lock Duty Cycle to 100%, Capillary Voltage 1600V, Dry Gas 3 l/min, Dry Temp 180°C, PASEF settings: 10 MS/MS scans (total cycle time 1.27sec), charge range 0-5, active exclusion for 0.4 min, Scheduling Target intensity 10000, Intensity threshold 2500, CID collision energy 42eV.

### TGF-beta1 Quantification

The serum level of TGF-β1 was measured using a commercially available Bio-Plex Pro TGF-β Assay (Bio-Rad Laboratories Ltd, Hercules, CA, USA) on the Bio-Plex™ 200 System. The Bio-Plex™ 200 software version 5.1.1 (Bio-Rad Laboratories) was used to determine the concentration in pg/mL.

## QUANTIFICATION AND STATISTICAL ANALYSIS

### Data processing and statistical analysis for mass spectrometry-based proteomics

Raw data files were analysed by Fragpipe (v15.0) using a protein sequence database of reviewed Murine proteins, accessed on 13/01/2021 from UniProt. The database contains 34278 entries including decoys which were generated and appended to the original using MSFragger (v3.2). Tryptic cleavage specificity was applied allowing for 2 missed cleavages, alongside fixed carbamidomethyl cysteine and variable methionine oxidation and N-terminal acetylation modifications. The peptide length and mass ranges were set to 7-50 residues and 500-5000 Da, respectively. The precursor mass error was set to -20-20 ppm and the fragment mass error was set to 10 ppm with mass calibration and parameter optimization enabled. The peptide spectrum matches were filtered using PeptideProphet and ProteinProphet in Philosopher (v3.4.13) to 1% PSM and 1% FDR. The peptides and proteins were quantified using IonQuant (v1.5.5) with the m/z, retention time and ion mobility tolerances set to 10 ppm, 0.4 minutes, 0.05 1/k0, respectively. The protein quant was performed using the MaxLFQ algorithm and proteins needed a minimum of 2 peptides to be quantified.

Data processing and analysis were performed using R (version 4.0.4). Only proteins that were quantified in at least 50% of replicates in at least one condition were kept. The protein intensities were log_2_-transformed. Missing values were imputed by using Missing Not At Random (MNAR) method implemented in msImpute R-package(v.1.2.0). The data were normalized using RUVSeq (Risso et al., 2014). The optimum k value used to remove the unwanted variation was determined based on PCA, RLE and p-value distribution plots. The R-package DEqMS (v. 1.9.0) was used to perform the differential analysis. Unlike limma, DEqMS estimates different prior variances for proteins quantified by different number of peptides, therefore achieving better accuracy compared to limma (Zhu et al., 2020). A protein was determined to be significantly differentially expressed if the false discovery rate (FDR) adjusted p-value was□≤□0.05. R-packages; ggplot2 (v. 3.3.3) and superheat (v. 0.1.0) were used to visualise the results.

EMT signature genes were obtained from Simeonov *et a*l, 2020. Protein expression of EMT signature genes, where detected, were summarised using the ssGSEA algorithm as implemented in the GSVA package for R. The statistical significance of differences in the ssGSEA signature score between groups was calculated using the Welch’s t-test, and Benjamini-Hochberg correction performed, as implemented in the R statistical package.

### Analysis of publicly available gene expression datasets

TCGA pancreatic cancer z distribution-transformed expression data was obtained from cbioportal.org (PMID 23550210) with correlating histopathological classifications (PMID 28810144, (Integrated Genomic Characterization of Pancreatic Ductal Adenocarcinoma, 2017). Gene expression differences between groups were ranked based on difference in mean z-score. For gene set enrichment analysis of TCGA gene expression data, the ssGSEA algorithm was used as implemented in the GSVA package for R. Patient survival analysis was performed using a publicly available webtool (http://www.oncolnc.org/) (Anaya, 2016). Gene expression comparison between human PDAC from TCGA and normal pancreas from GTEX were performed using Gene Expression Profiling Interactive Analysis (GEPIA) webtool (http://gepia.cancer-pku.cn/) (Tang *et al*., 2017).

### General statistical analysis

Organoid morphology quantification of either glandular of solid were manually quantified using ImageJ.

All results were plotted in GraphPad Prism. The statistical test performed has been described in the legend of each figure. A result was considered significant if the P-value was less than 0.05 and has been indicated in graphs as * P-value <0.05, ** P-value <0.01, *** P-value <0.001 and *** P-value <0.0001.

