## Supplementary figures for "TGF-β1 induced S100 family protein expression is associated with epithelial to mesenchymal transition states and poor survival in pancreatic cancer"

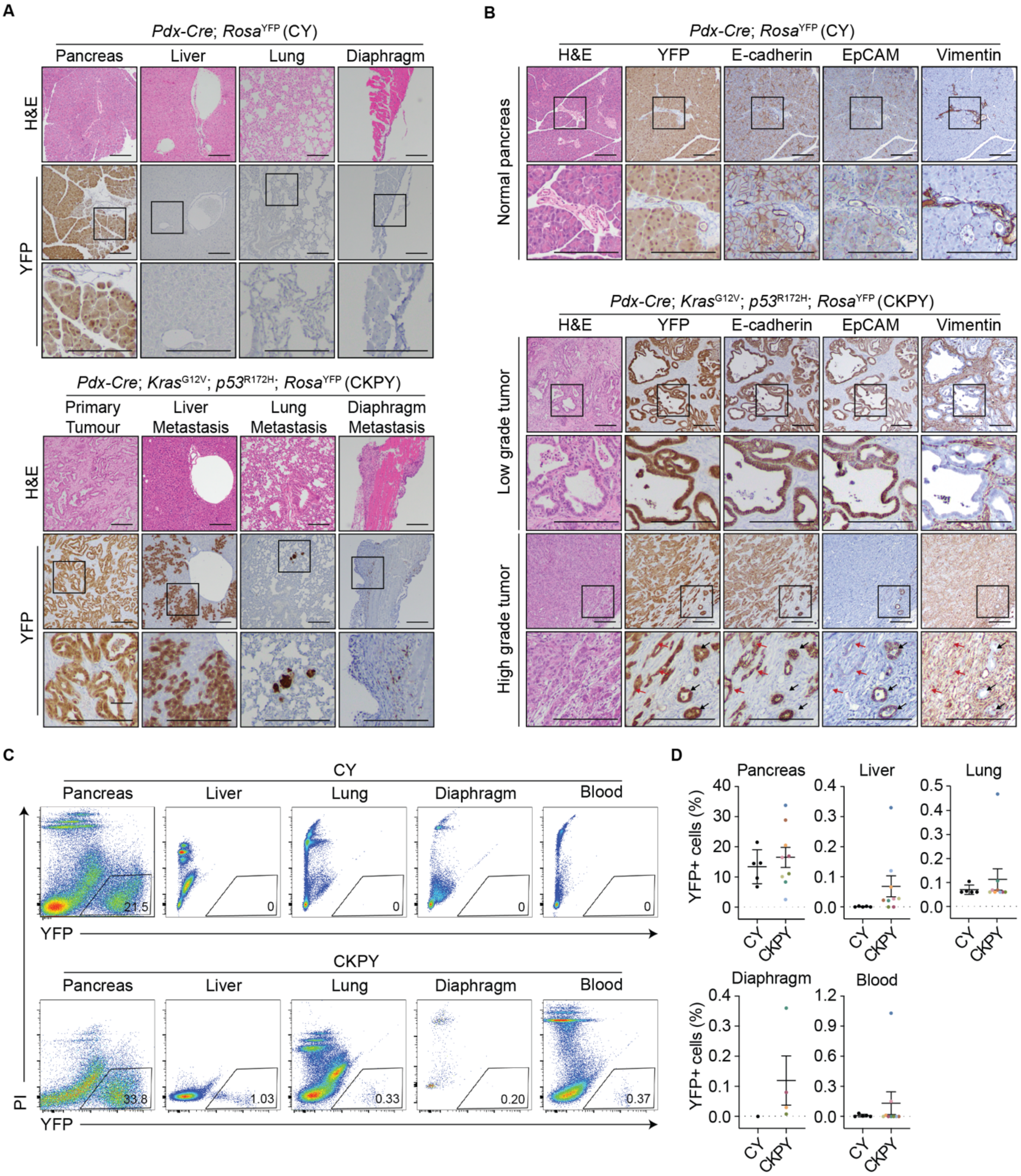
A
Pdx-Cre; Rosa YFP (CY)
Pancreas
Liver
Lung
Diaphragm
H&E
YFP
Pdx-Cre; Kras G12V; p53 R172H; Rosa YFP (CKPY)
Primary Tumour
Liver Metastasis
Lung Metastasis
Diaphragm Metastasis
B
Pdx-Cre; Rosa YFP (CY)
H&E
YFP
E-cadherin
EpCAM
Vimentin
Normal pancreas
Pdx-Cre; Kras G12V; p53 R172H; Rosa YFP (CKPY)
Low grade tumor
High grade tumor
C
CY
Pancreas
Liver
Lung
Diaphragm
Blood
21.5
0
0
0
0
YFP
CKPY
33.8
1.03
0.33
0.20
0.37
PI
YFP
D
Pancreas
Liver
Lung
Diaphragm
Blood
YFP+ cells (%)
CY
CKPY

**Fig S1. Identification and isolation of YFP+ cells from secondary tumors (related to Figure 1)**

(A) Representative images of the pancreas, liver, lung and diaphragm of CY mice (N=5) and CKPY (N=9) mice. Adjacent images of H&E and YFP immunohistochemsitry are shown. Scale bar = 200 μm.

(B) Representative images of normal pancreas from CY mice (top) and primary tumors from CKPY mice, separated by low grade tumor (middle) and high grade tumor (bottom). Adjacent H&E and YFP, E-cadherin, EpCAM and Vimentin stained sections are shown. YFP+EpCAM+ (black arrow) and YFP+EpCAM- (red arrow) cells are present in the high grade tumor. Scale bar = 200 μm.

(C) FACS gating strategy for YFP+ cell sorting from the pancreas, liver, lung, diaphragm and blood harvested from CY mice (N=5) and CKPY mice (N=9).

(D) Percentage of live YFP+ cells present in the indicated organs or blood. Each dot represents an individual mouse. Data is presented as mean +/- SEM.

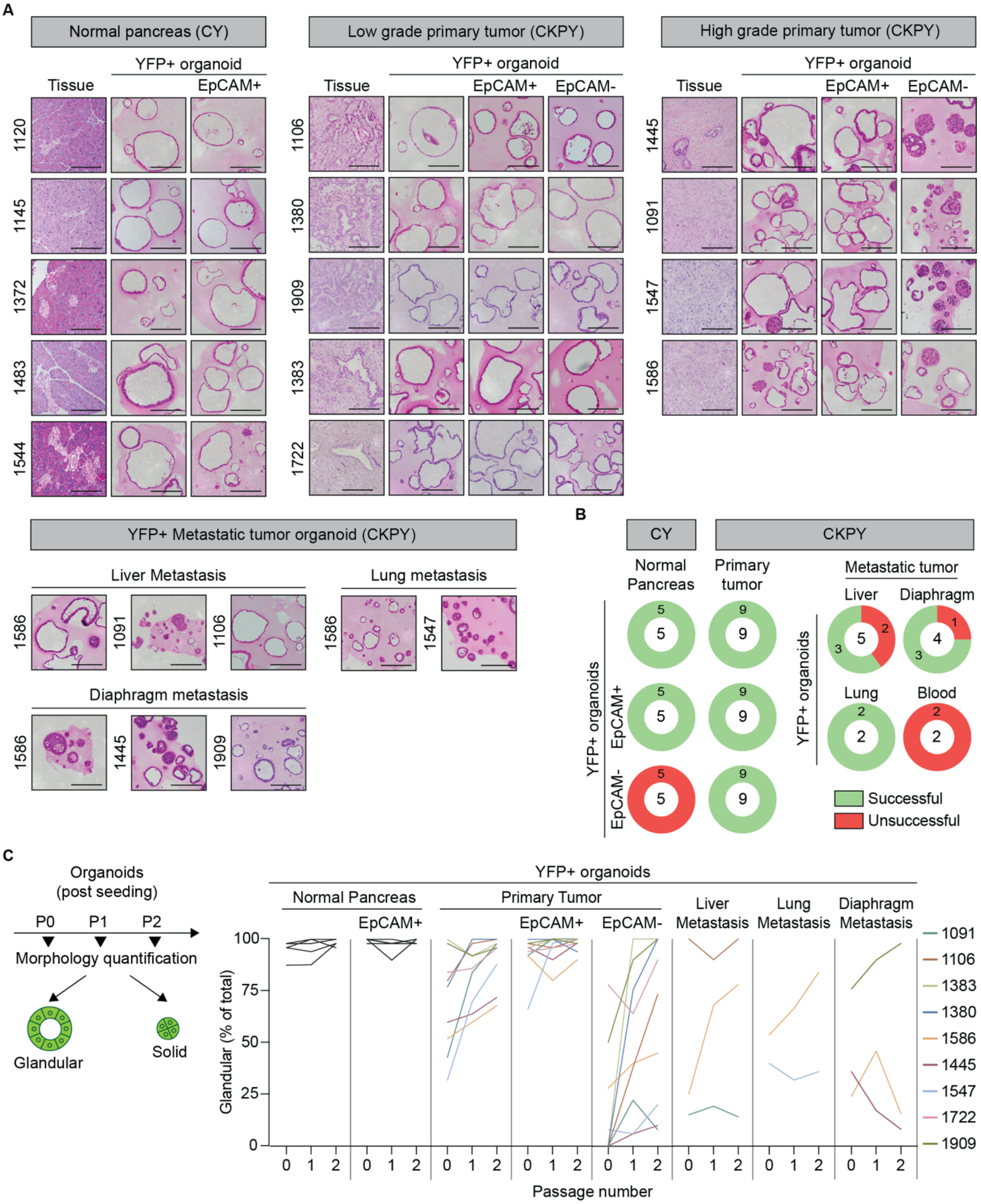
A
Normal pancreas (CY)
YFP+ organoid
Tissue
EpCAM+
1120
1145
1372
1483
1544
Low grade primary tumor (CKPY)
YFP+ organoid
Tissue
EpCAM+
EpCAM-
1106
1380
1909
1383
1722
High grade primary tumor (CKPY)
YFP+ organoid
Tissue
EpCAM+
EpCAM-
1445
1091
1547
1586
YFP+ Metastatic tumor organoid (CKPY)
Liver Metastasis
1586
1091
1106
Lung metastasis
1586
1547
Diaphragm metastasis
1586
1445
1909
B
CY
CKPY
Normal Pancreas
Primary tumor
Metastatic tumor
Liver
Diaphragm
Lung
Blood
YFP+ organoids
EpCAM+
EpCAM-
Successful
Unsuccessful
C
Organoids (post seeding)
P0
P1
P2
Morphology quantification
Glandular
Solid
YFP+ organoids
Normal Pancreas
Primary Tumor
EpCAM+
EpCAM-
Liver Metastasis
Lung Metastasis
Diaphragm Metastasis
Glandular (% of total)
Passage number
1091
1106
1383
1380
1586
1445
1547
1722
1909

**Fig S2. Successful growth of murine organoids from multiple tissues (related to Figure 1)**

(A) Representative H&E images of the primary tissue and corresponding normal pancreatic organoids generated from CY mice (N=5), or tumor organoids generated from CKPY mice (N=5 low grade, N= 4 high grade). Scale bar = 100µm.

(B) Number of organoid lines generated from live YFP+ cells sorted from the indicated organs of CY mice (N=5) or CKPY mice (N=9). Center of the plot indicates total number of organs with sorted YFP+ cells for organoid generation. Scale bar = 100µm.

(C) Quantification of the stability of organoid morphology following serial passaging (P) post seeding. Presented as the percent of glandular organoids present within the culture following P0, P1 P2 for normal (N=5), primary tumor (N=9) and metastatic tumor organoids (N=8). Data is presented as relative to total number of organoids within each well. Individual organoid lines are indicated.

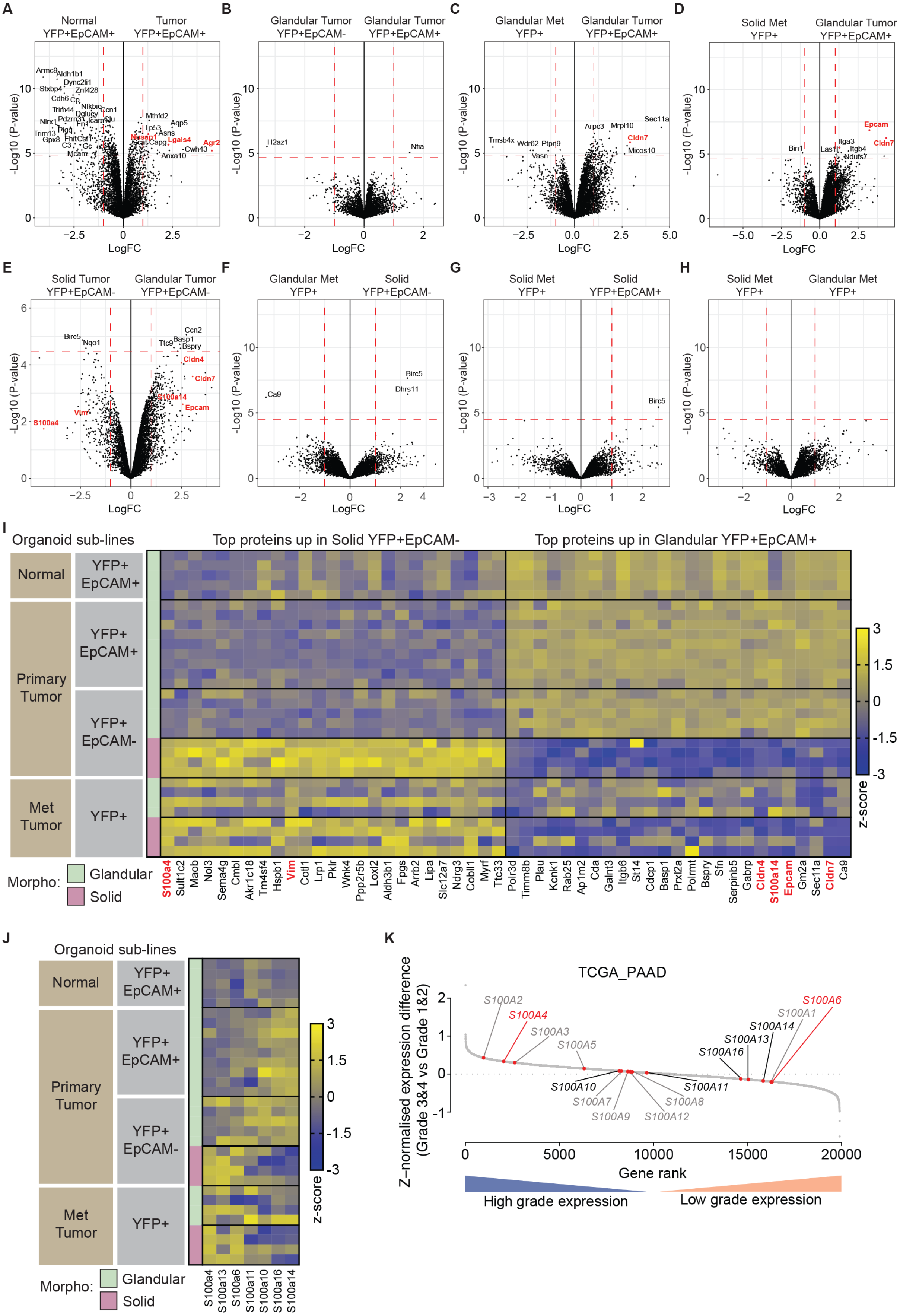
A
Normal YFP+EpCAM+
Tumor YFP+EpCAM+
B
Glandular Tumor YFP+EpCAM-
Glandular Tumor YFP+EpCAM+
C
Glandular Met YFP+
Glandular Tumor YFP+EpCAM+
D
Solid Met YFP+
Glandular Tumor YFP+EpCAM+
E
Solid Tumor YFP+EpCAM-
Glandular Tumor YFP+EpCAM-
F
Glandular Met YFP+
Solid YFP+EpCAM-
G
Solid Met YFP+
Solid YFP+EpCAM+
H
Solid Met YFP+
Glandular Met YFP+
-Log10 (P-value)
LogFC
I
Organoid sub-lines
Top proteins up in Solid YFP+EpCAM-
Top proteins up in Glandular YFP+EpCAM+
Normal
Primary Tumor
Met Tumor
YFP+ EpCAM+
YFP+ EpCAM-
YFP+
Morpho: Glandular Solid
z-score
S100a4 Sult1c2 Maob Nol3 Sema4g Cmbl Akr1c18 Tm4sf4 Hspb1 Vim Cotl1 Lrp1 Pklr Wnk4 Ppp2r5b Loxl2 Aldh3b1 Fpgs Arb2 Lipa Slc12a7 Ndrg3 Cobll1 Myrf Ttc33 Polr3d Timm8b Plau Kcnk1 Rab25 Ap1m2 Cda Galnt3 Itgb6 St14 Cdcp1 Basp1 Prxl2a Polrmt Bspry Sfn Serpinb5 Gabrp Cldn4 S100a14 Epcam Gm2a Sec11a Cldn7 Ca9
J
Organoid sub-lines
Normal
Primary Tumor
Met Tumor
Morpho: Glandular Solid
z-score
S100a4 S100a13 S100a6 S100a11 S100a10 S100a16 S100a14
K
TCGA_PAAD
Z-normalised expression difference (Grade 3&4 vs Grade 1&2)
Gene rank
High grade expression
Low grade expression
S100A2 S100A4 S100A3 S100A5 S100A10 S100A7 S100A9 S100A12 S100A8 S100A11 S100A16 S100A13 S100A14 S100A1 S100A6

**Fig S3. Proteomics analysis reveals different expression profile in the murine organoid sub-lines (related to Figure 2)**

(A-H) Volcano plot illustrating the $log_2$ protein ratios in whole cell lysates of MO, comparing glandular YFP+EpCAM+ tumor organoid with normal YFP+EpCAM+ organoid (A); glandular YFP+EpCAM- tumor organoid (B); glandular YFP+ metastatic organoid (C); and solid YFP+ metastatic organoid (D); YFP+EpCAM- tumor organoids (E); metastatic organoids (F-H). Proteins were deemed differentially regulated if the log2 fold change in protein expression was $\geq$ 1-fold and exhibited an adjusted *p*-value $\leq$ 0.05.

(I-J) Heat-map representation of the top 25 differentially expressed proteins between glandular YFP+EpCAM+ tumor organoid and solid YFP+EpCAM- tumor organoid (I); S100a family proteins (J) in all the organoid lines utilized for proteomcis analysis including normal YFP+EpCAM+ (N=5), tumor YFP+EpCAM+ (N=9), tumor YFP+EpCAM- (N=9) and YFP+ metastatic (N=8) tumor organoids. Organoids are grouped as glandular (G) and solid (S) morphologies.

(K) Expression of S100A family members in patient tumors ranked by the mean expression difference in grade 1/2 tumor v.s. grade 3/4 tumor from high purity tumor (Raphael *et al.*, 2017) from the TCGA-PAAD dataset. S100 family members detected in proteomics (black arrow), protein of interest (red) and other S100 family members (grey).

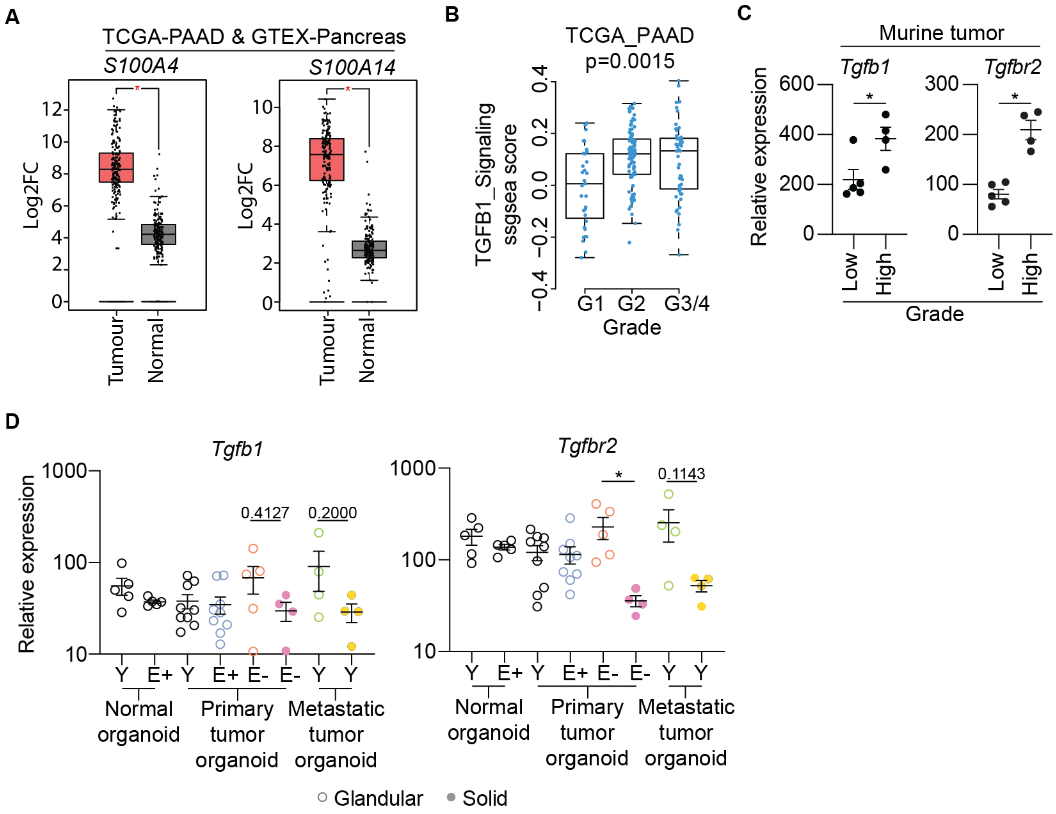


**Fig S4. Murine organoids express *Tgfb1* and *Tgfbr2* (related to Figure 3)**

(A) RNA expression analysis of *S100A4* and *S100A14* in human PDAC (TCGA-PAAD, n = 179) and normal pancreas (GTEX-pancreas, n = 171) from the GEPIA tool (Tang *et al.*, 2017).

(B) GSEA plot evaluating the TGF-beta signaling signature (KARAKAS_TGFB1_signaling) between grade 1, grade 2 and grade ¾ patients tumor from the TCGA-PAAD dataset.

(C) mRNA expression levels of *Tgfb1* and *Tgfbr2* in the original murine tumor. Each dot represents an individual tumor. Each dot represents an individual mouse. Data is relative to *Gapdh*, presented as mean +/- SEM. . *p<0.05, Mann-Whitney test.

(D) mRNA expression of *Tgfb1* and *Tgfbr2* in YFP+ (Y), YFP+EpCAM+ (E+) and YFP+EpCAM- (E-) organoids. Each dot represents an individual MO. Data is relative to *Gapdh*, presented as mean +/- SEM. *p<0.05, Mann-Whitney test.

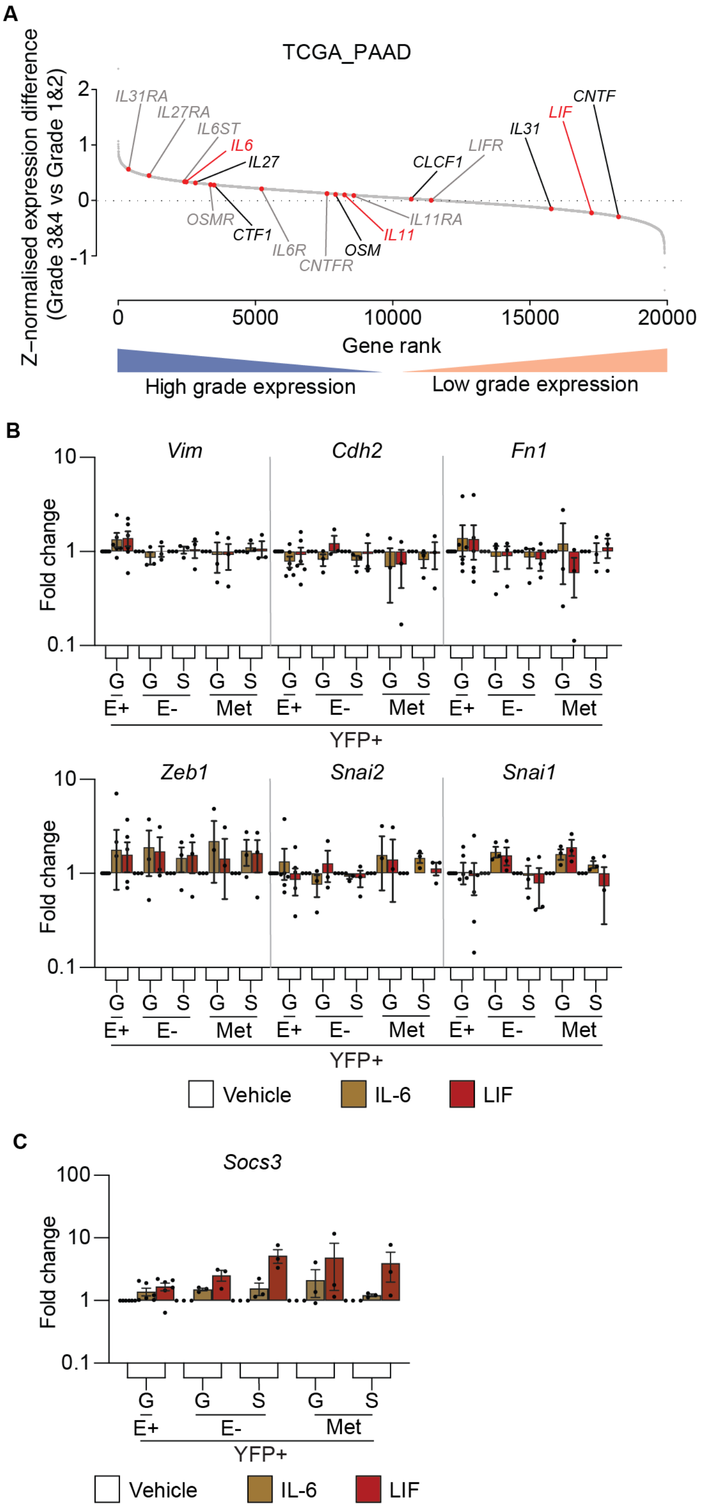

A
TCGA_PAAD
Z-normalised expression difference (Grade 3&4 vs Grade 1&2)
IL31RA
IL27RA
IL6ST
IL6
IL27
OSMR
CTF1
IL6R
CNTFR
OSM
IL11
IL11RA
CLCF1
LIFR
IL31
LIF
CNTF
Gene rank
High grade expression
Low grade expression
B
Vim
Cdh2
Fn1
Zeb1
Snai2
Snai1
Fold change
G
S
E+
E-
Met
YFP+
Vehicle
IL-6
LIF
C
Socs3
Fold change
G
S
E+
E-
Met
YFP+
Vehicle
IL-6
LIF

**Figure S5. IL-6 or LIF, do not alter tumor organoid morphology or mesenchymal signature (related to Figure 5)**

(A) Expression of IL-6 family members in patients tumor ranked by the mean expression difference in grade 1/2 tumor v.s. grade 3/4 tumor from all tumors from the TCGA-PAAD dataset. Cytokines (black arrow), cytokines of interest (red) and cytokine receptors (grey).

(B-C) mRNA expression levels of mesenchymal markers, *Vim*, *Cdh2*, *Fn1*, *Snai1*, *Snai2*, *Zeb1* (B); and the IL-6 cytokine family target gene, *Socs3* (C), in YFP+EpCAM+ (E+), YFP+EpCAM- (E-) primary tumor organoids and YFP+ metastatic organoids following the addition of the indicated cytokine (IL-6 brown, LIF red). Each dot represents an individual organoid. Organoids are grouped as glandular (G) and solid (S) morphologies. Data includes 3 biological replicates and is presented as log10 fold change relative to the vehicle (white) control, mean +/- SEM. paired t-test.

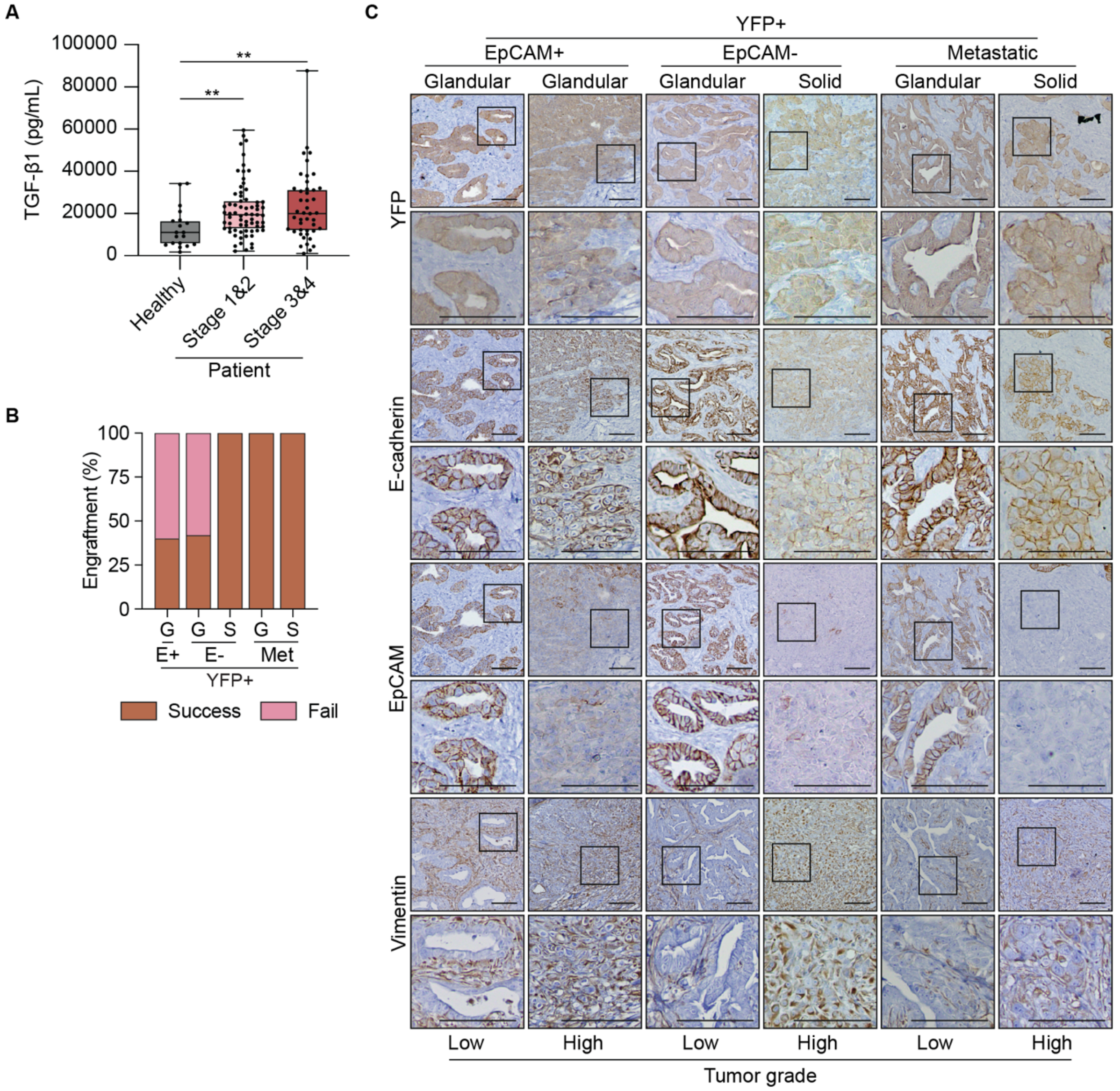


**Fig S6. Generation of a murine allograft biobank with different histological features (related to Figure 6)**

(A) Serum TGFβ1 in healthy individuals (n = 19) compared to PDAC patients grouped by stage 1/2 (n = 69) and stage 3/4 (n = 45). Data is presented as box and whisker plot. *p<0.05, *p<0.01, Mann-Whitney test.

(B) Successful generation of allografts from YFP+EpCAM+ (E+), YFP+EpCAM- (E-) primary tumor organoids and YFP+ metastatic organoids. Organoids are grouped as glandular (G) and solid (S) morphologies. Each bar graph indicates the overall number of allografts generated by subcutaneous transplantation (N= 9 mice per organoid line).

(C) Representative immunohistochemical staining of allografts generated from YFP+EpCAM+ (E+), YFP+EpCAM- (E-) primary tumor organoids and YFP+ metastatic organoids for YFP, E-Cadherin, and EpCAM. Organoids are grouped as glandular (G) and solid (S) morphologies.

Scale bar = 200µm.
